# Tnfrsf10 Signaling is Required to Maintain the Stem Cell Niche in the Zebrafish Lateral Line

**DOI:** 10.1101/2025.04.18.649014

**Authors:** Chunxin Fan, Xiaoyi Yuan, Jian Wang, Wuhong Pei, Hongfei Zheng, Xinyu Zhang, Shawn M. Burgess

## Abstract

Regenerative capacity varies significantly among species and across different organs. In mammals, the inner ear exhibits a limited ability to regenerate hair cells, whereas the lateral line sensory organ of zebrafish demonstrates apparently limitless capability to spontaneously regenerate hair cells following injury. However, the specific cell populations representing the stem cells in the neuromast and the mechanism of stem cell self-renewal remain unclear. In this study, we show that a mutation in *tnfrsfa* led to the depletion of neuromast sensory organs and impairment of hair cell regeneration in zebrafish. The decrease of cells peripheral to the neuromasts in *tnfrsfa* mutants indicates that these peripheral cells, particularly cells called “mantle cells”, serve as the stem cells within the neuromast. Furthermore, our findings indicate that the ligands Tnfsf10 and Tnfsf10l3 (TRAIL) act through Tnfrsfa activating the NF-κB signaling pathway in the mantle cells in a non-cell-autonomous manner. NF-κB signaling is crucial for maintaining the stem cell identity of the mantle cells through activation or maintenance of *sox2* expression. These findings demonstrate a critical role for TNF superfamily signaling in stem cell maintenance.

**Highlights:**

- Zebrafish *tnfrsfa* maintains the stem cell niche in neuromasts and is necessary for hair cell regeneration of the lateral line
- Tnfrsfa is essential for mantle cell proliferation during lateral line development and hair cell regeneration
- Tnfsf10 and Tnfsf10l3 signal through Tnfrsfa, activating NF-κB signaling and Sox2 expression in the mantle cells in a non-cell-autonomous manner
- Tnfrsfa regulates the development of neuromast independently of the Notch signaling pathway

## Introduction

Regenerative capacity shows considerable variation among species and across different organs. The mechanosensory receptor in the inner ear of all vertebrates is the hair cell and the mammalian inner ear possesses severely-limited hair cell regenerative capacity, and only at very young ages (Burns et al., 2012; Cox et al., 2014). Sensorineural deafness caused by hair cell loss either through age or environmental exposures is the main reason for human hearing loss. However, the hair cells in the non-mammalian inner ear or the lateral line sensory organs of fish and amphibians can regenerate spontaneously after injury (Corwin and Cotanche, 1988; Ryals and Rubel, 1988). The regenerative potential of the inner ear or lateral line organ is dependent on the availability of stem cells or progenitors that are able to functionally replace the lost or damaged hair cells (Ghysen and Dambly-Chaudière, 2007; Kelley and Meyers, 2019). Pinpointing the origin and regulation of stem cells in regenerating species is of significant biological and clinical interest.

The lateral line neuromast is an ideal model for studying hair cell regeneration due to its location on the skin giving it experimental accessibility. The neuromast is comprised of centrally located sensory hair cells, underlying non-sensory “support cells”, and a peripheral ring of “mantle cells” (Ghysen and Dambly-Chaudière, 2007; Ma and Raible, 2009). Despite similarities in morphology, support cells can be classified into anterior-posterior (A/P), dorsoventral (D/V), and central subpopulations based on lineage tracing and single-cell RNA sequencing (scRNA-seq) data from regenerating neuromasts. Central support cells act as immediate hair cell precursors, replenishing hair cells through mitotic division and differentiation. D/V support cells undergo symmetrical division, generating two new support cells, whereas A/P support cells remain relatively inactive under normal conditions, activating only to divide and differentiate following severe injury (Denans et al., 2019; Romero-Carvajal et al., 2015; Thomas and Raible, 2019; Viader-Llargués et al., 2018; Wibowo et al., 2011). Similarly, mantle cells can proliferate and differentiate into all neuromast cell types (Dufourcq et al., 2006; Seleit et al., 2017; Williams and Holder, 2000). However, the relationship of mantle cells and support cells to the stem cell niche of the neuromasts has yet to be fully established (Thomas and Raible, 2019).

Pro-inflammatory factors play a crucial role in cell proliferation and survival, essential for regeneration of various tissues including the spinal cord, limbs, fins, retina, liver, and heart (Anguita-Salinas et al., 2019; Cooke, 2019; Hasegawa et al., 2017; Karin and Clevers, 2016; Kizil et al., 2012; Lai et al., 2019; Li et al., 2012; Tsarouchas et al., 2018). The role of inflammatory signaling pathways in tissue regeneration is conserved across different vertebrates, including zebrafish, geckos, and mice (He et al., 2015; Iribarne, 2021; Zhang et al., 2024a). Recent single-cell RNA sequencing (scRNA-seq) studies of regenerating neuromasts have revealed that inflammatory factors are rapidly upregulated following hair cell ablation, underscoring their pivotal role in hair cell regeneration (Baek et al., 2022).

Tumor necrosis factor superfamily (TNFSF) members are key pro-inflammatory factors. These factors function through the trimerization of soluble or membrane-bound TNF ligands, which interact with corresponding trimerized tumor necrosis factor receptor superfamily (TNFRSF) members. This interaction occurs between the TNF homology domain (THD) of the ligands and the cysteine-rich domain (CRD) of the receptors. TNFSF ligands and TNFRSF receptors play critical roles in cell apoptosis, survival, proliferation, and differentiation by activating downstream signaling pathways, including Caspase, NF-κB, and JNK (Dostert et al., 2019). Recent research has highlighted the involvement of various TNFSF members and the downstream NF-κB signaling pathway in stem cell emergence and reprogramming (Campbell et al., 2024; Espín-Palazón et al., 2014; He et al., 2015; Nelson et al., 2013; Palazzo et al., 2020; Zhang et al., 2024b).

In this study, we generated *tnfrsfa* mutant zebrafish lines and demonstrated that Tnfrsfa is essential for both the growth of the lateral line sensory organ and the regeneration of hair cells. Specifically, mutations in *tnfrsfa* lead to the decrease in the number of peripheral cells and defects in the proliferation of mantle cells during hair cell regeneration. In addition, hair cell regeneration was impacted but only when repeated ablations occurred, indicating that signaling through Tnfrsfa is essential for maintaining the stem cell niche within the neuromast but not the regeneration process specifically. Additionally, we found that the ligands for Tnfrsfa, Tnfsf10 and Tnfsf10l3, activate the NF-κB signaling pathway and Sox2 in the mantle cells. The role of Tnfrsfa in neuromast development was independent of the Notch signaling pathway which is a known driver of hair cell differentiation and regeneration. This research identifies a critical pro-inflammatory signaling pathway essential for maintaining the stem cell niche necessary for hair cell regeneration and presents a potential target for promoting mammalian hair cell regeneration.

## Results

### Impact of tnfrsfa mutations on lateral line sensory organ and hair cell regeneration

Previous studies showed strong expression of zebrafish *tnfrsfa* (also known as *zOTR*) in the lateral line neuromast (Eimon et al., 2006) and we identified *tnfrsfa* as a candidate regeneration gene in a targeted genetic screen (Pei et al., 2018). We confirmed *tnfrsfa* expression in the posterior lateral line primordium at 24 hours post-fertilization (hpf) and in the neuromast at 48 hpf using in situ hybridization (Fig. 1A). To explore the role of the *tnfrsfa* gene in lateral line development and hair cell regeneration, we established two independent zebrafish *tnfrsfa* mutant lines using CRISPR/Cas9, *tnfrsfa^crd^* and *tnfrsfa^dd^*, frame-shift mutations that occur upstream of the first cysteine-rich domain (CRD1) or the death domain (DD) respectively. The *tnfrsfa^crd^* mutation involves a four-nucleotide (GTCA) deletion in exon 3, predicted to produce truncated proteins with a partial CRD1 domain (Fig. 1B). qRT-PCR analysis confirmed a significant reduction in *tnfrsfa* mRNA in *tnfrsfa^crd^* homozygous mutants, indicating nonsense-mediated decay of the mutant transcripts (Fig. 1C). *tnfrsfa^crd^* homozygous mutants are viable with no obvious phenotypic changes, except for reduced fertility. Analysis of hair cell dynamics in the posterior lateral line neuromasts (P1-P4) using vital dye YO-PRO-1 revealed slightly more hair cells at 72 hpf (p<0.01) but fewer by 5 dpf (p<0.001) in *tnfrsfa^crd/crd^* mutants compared to that in wild-type siblings (Fig. 1D). These differences in hair cell numbers were confirmed by Otoferlin (HCS-1) antibody immunostaining (Fig. 1E). The *tnfrsfa^dd^*mutants produce a protein with an incomplete DD. Both *tnfrsfa^dd^*homozygous and heterozygous mutants exhibited phenotypes similar to *tnfrsfa^crd/crd^*mutants with a significant reduction in neuromast hair cells by 5 dpf, suggesting a dominant negative effect of the *tnfrsfa^dd^*mutation (Supplementary Fig. S1).

**Figure 1.**
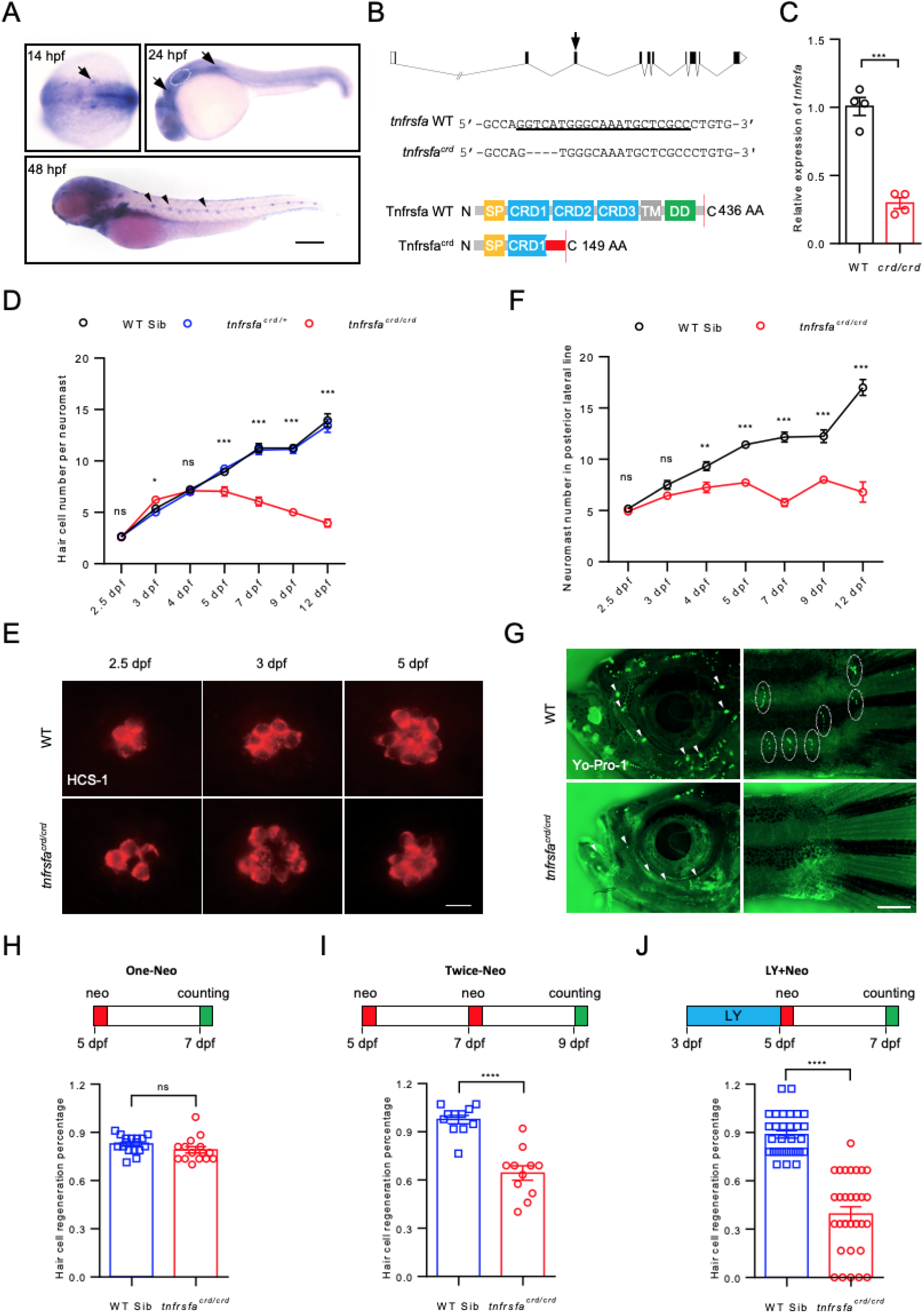
Impact of *tnfrsfa* Deficiency on Lateral Line Development and Hair Cell Regeneration. (A) Whole-mount *in situ* hybridization showing *tnfrsfa* expression in the lateral line primordium (arrow), neuromasts (arrowhead), and otic vesicle (dashed circle) in zebrafish embryos. Scale bar = 500 μm. (B) Schematic of *tnfrsfa* mutations generated using CRISPR/Cas9. Upper panel: gRNA location (arrow) within the *tnfrsfa* locus. Middle panel: sequences surrounding the gRNA target in wild-type and mutant *tnfrsfa* alleles (gRNA target underlined). Bottom panel: predicted protein structure of wild-type and mutant Tnfrsfa. SP, signal peptide; CRD, cysteine-rich domain; TM, transmembrane domain; DD, death domain. Red box represents the missense sequence before the stop codon. (C) qRT-PCR analysis of *tnfrsfa* expression in wild-type and *tnfrsfa^crd/crd^* embryos at 48 hpf, mRNA reduced through nonsense mediated decay. N=4; ***, p < 0.001. (D) Quantification of hair cells in the posterior lateral line using Yo-Pro-1 staining in wild-type and *tnfrsfa^crd/crd^*mutant larvae during development. Mutant embryos had more hair cells on average at 3 dpf and dropped off after that. N = 5-30; *, p < 0.05; ***, p < 0.001; ns, not significant. E) Immunostaining of neuromast hair cells in wild-type and *tnfrsfa^crd/crd^* mutants using the HCS1 antibody labeling mature hair cells. Scale bar = 10 μm. (F) Quantification of total posterior lateral line neuromasts in wild-type and *tnfrsfa^crd/crd^* mutants. N = 5-17; **, p < 0.01; ***, p < 0.001; ns, not significant. (G) Visualization of superficial (arrowhead) and canal neuromasts (dashed circle) in wild-type and *tnfrsfa^crd/crd^*mutant adults at 3 months post-fertilization (mpf) using Yo-Pro-1 staining. Mutant adults had almost no visible neuromasts. (H-J) Quantification of hair cell regeneration in the posterior lateral line of wild-type and *tnfrsfa^crd/crd^*mutant larvae after after single a neomycin treatment (One-Neo, N=15-17) (H), two sequential neomycin treatments (Twice-Neo, N=11) (I), neomycin treatment following LY411575 incubation from 3 dpf to 5 dpf (LY and Neo, N=30) (J). ***, p < 0.001; ****, p < 0.0001.

In addition to affecting hair cell number, *tnfrsfa^crd^*mutations also led to a decrease in the number of neuromasts at larval stages. By 4 dpf, *tnfrsfa^crd^* mutants had fewer posterior lateral line neuromasts compared to wild-type siblings, and the difference in neuromast number increased between *tnfrsfa^crd/crd^* mutants and wild-type siblings gradually (Fig. 1F). Notably, the superficial neuromasts were completely absent and the cranial canal neuromasts were significantly smaller in adult *tnfrsfa^crd/crd^* mutants (3 months post-fertilization, mpf) (Fig. 1G). To examine if the mutation also impacted inner ear hair cells, we dissected saccules from adult *tnfrsfa^crd/crd^* mutant and wild-type siblings and found that there was no significant reduction in the saccule hair cells in *tnfrsfa^crd/crd^*mutant adults compared with wild-type siblings (Supplementary Fig. S2). These results indicate that *tnfrsfa* is essential for the maintenance of lateral line sensory organs in zebrafish but not inner ear homeostasis.

Lateral line hair cells regenerate robustly after ablation with aminoglycoside antibiotics or copper sulfate (Behra et al., 2009; Cruz et al., 2015; Harris et al., 2003; Song et al., 1995; Williams and Holder, 2000). To investigate the role of the *tnfrsfa* gene in hair cell regeneration, we ablated hair cells by treating larvae at 5 dpf with neomycin (One-Neo) and evaluated the hair cell regeneration after two days-recovery.

Results showed that the total hair cell number was down in *tnfrsfa^crd/crd^* mutants before ablation but the rate of return to the previous hair cell numbers were comparable to that in wild-type siblings, indicating *tnfrsfa* does not significantly influence hair cell regeneration following a single neomycin treatment (Fig. 1H). We performed repeated neuromast injury by treating zebrafish larvae with neomycin at 5 dpf, followed by another neomycin treatment at 7 dpf (Twice-Neo). Upon the repeated injury, hair cell recovery decreased significantly in *tnfrsfa^crd/crd^* mutants compared to that in wild-type siblings (Fig. 1I). LY411575, a Notch signaling inhibitor, transforms support cells into supernumerary hair cells through release of lateral inhibition as described previously (Romero-Carvajal et al., 2015; Ye et al., 2020) “using up” the available support cells. The mutants showed hair cell regeneration defects when pretreated with LY411575 for 2 days followed by a neomycin treatment at 5 dpf (LY+Neo) similar to the double treatment with neomycin (Fig. 1J), while in wild-type fish, the neuromasts were able to generate new support cells and hair cells after ablation. These results indicate that *tnfrsfa* is required for maintaining hair cell regeneration over longer time periods in zebrafish but is not an absolute requirement for support cells to differentiate into new hair cells after mild hair cell injuries.

### tnfrsfa maintains accessory cells in a cell-non-autonomous manner

Accessory cells, including mantle cells and support cells, serve as stem/progenitor cells within neuromasts (Dufourcq et al., 2006; Romero-Carvajal et al., 2015; Seleit et al., 2017; Thomas and Raible, 2019). Crossing *tnfrsfa^crd/crd^* mutants with the *Tg(tnks1bp1:EGFP)* transgenic reporter line (hereafter referred to as *SCM1:EGFP*), which marks all accessory cells (Behra et al., 2012), showed that the shape of neuromasts in *tnfrsfa^crd/crd^*mutants was not altered, but the size of neuromasts in *tnfrsfa^crd/crd^* mutants was modestly but statistically significantly reduced (Fig. 2A-B). Immunostaining for Sox2, a marker for both support cells and mantle cells (Hernández et al., 2007), revealed that the number of Sox2-positive cells decreased significantly in *tnfrsfa^crd/crd^*mutants compared to wild-type siblings (Fig. 2C-D). In the Sox2 antibody staining, we also noticed wild-type fish had slightly weaker Sox2-positive cells in the periphery of the neuromasts and in some cells between the neuromasts (marked with white arrows in Fig. 2C). These cells appeared to be missing in *tnfrsfa^crd/crd^* mutant fish. In mature adults, essentially all the neuromasts were not detectable in *tnfrsfa^crd/crd^* mutants, as shown by an enhancer trap line that recapitulates *six2b* expression, *Tg(sou10:EGFP)* (*sou10:EGFP* for short) (Supplementary Fig. S3) (Fan et al., 2022).

**Figure 2.**
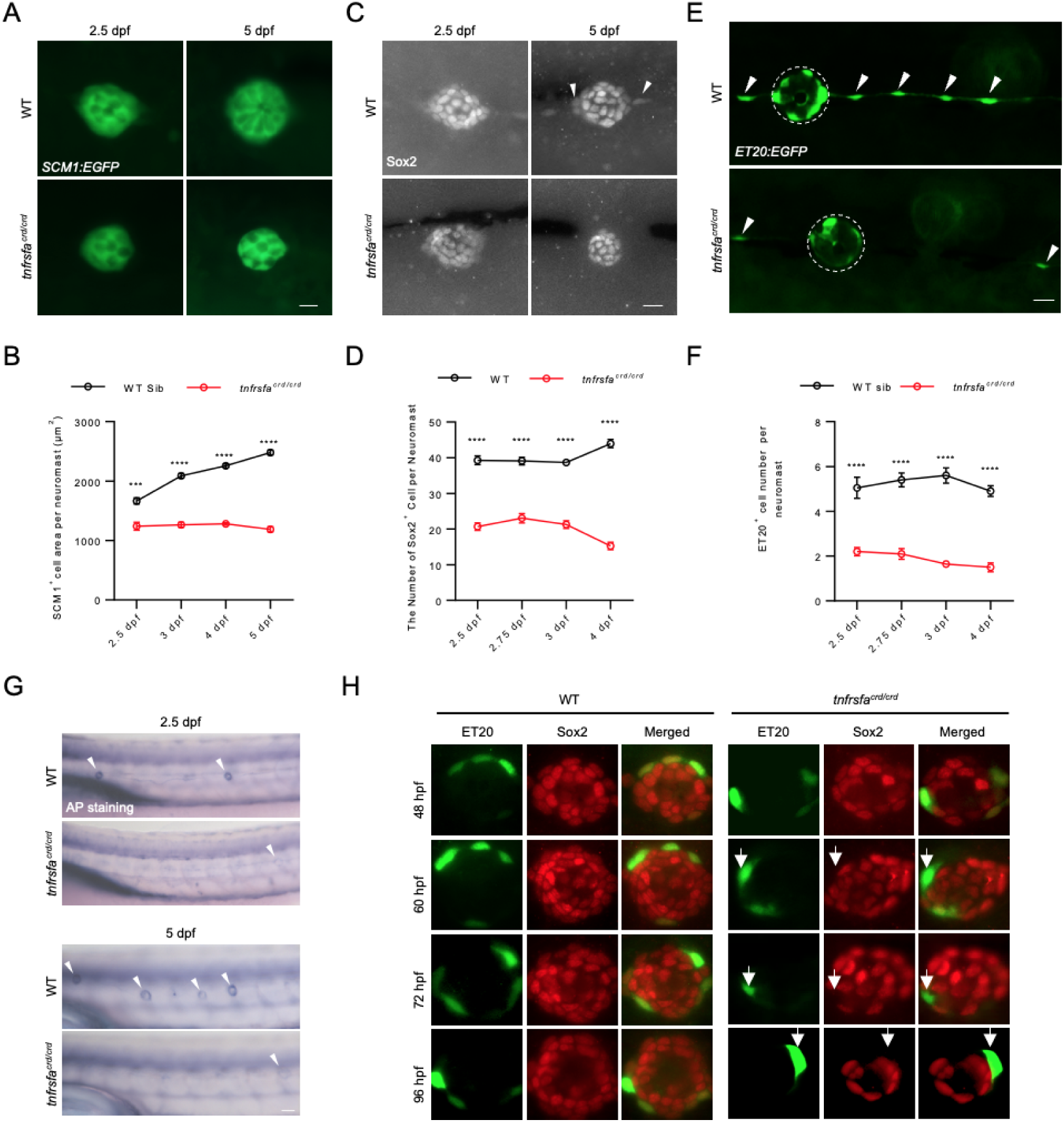
*tnfrsfa* is Required for Accessory Cell Maintenance in Neuromasts. (A) Live imaging of accessory cells in neuromasts in wild-type and *tnfrsfa* mutant larvae using the *Tg(SCM1:EGFP)* transgenic line. Scale bar = 10 μm. (B) Quantification of EGFP^+^ area in neuromasts of wild-type and *tnfrsfa* mutants using the *Tg(SCM1:EGFP)* transgenic line. N=8-16; ****, p < 0.0001. (C) Accessory cells visualized by immunostaining with Sox2 antibody in wild-type and *tnfrsfa^crd^*mutant siblings. Peripheral cells labeled by Sox2 in wild-types but missing in *tnfrsfa^crd^* mutant siblings marked by arrowheads. Scale bar, 10 μm. (D) Quantification of Sox2 positive cells in neuromasts in wild-type and tnfrsfacrd mutant siblings. N=20. ****, p < 0.0001. (E) Live imaging of mantle cells (dashed circle) and interneuromast cells (arrowhead) in wild-type and *tnfrsfa* mutant larvae using the *Tg(ET20:EGFP)* transgenic line. Mutant image shows rare positive cells, many fish had no positive cells. Scale bar = 10 μm. (F) Quantification of EGFP+ cells in neuromasts of wild-type and *tnfrsfa* mutants using *Tg*(*ET20:EGFP*). N=20; ****, p < 0.0001. (G) Alkaline phosphatase staining of the posterior lateral line in wild-type and *tnfrsfa* mutant larvae, with arrowheads indicating neuromasts. (H) Double immunostaining of *Tg*(*ET20:EGFP*) transgenic wild-type or *tnfrsfa* mutant embryos with anti-GFP (green) and anti-Sox2 (red) antibodies. Arrows indicate GFP positive mantle cells that are not expressing Sox2.

Using the *Et(krt4:EGFP)SqET20* transgenic line (hereafter referred to as *ET20:EGFP*) as a mantle cell reporter, we found that very few *tnfrsfa^crd/crd^* mutants contained *ET20:EGFP* positive cells within the neuromasts, and there was a significant reduction in the total number of mantle cells and inter-neuromast cells in the mutants compared to their wild-type siblings (Fig. 2E-F). Alkaline phosphatase (AP) is a marker for a variety of stem cells and has been used to label the neuromast (Lush et al., 2019; Štefková et al., 2015; Villablanca et al., 2006). The alkaline phosphatase activity is visible as a ring in wild-type neuromasts, while it was virtually undetectable in *tnfrsfa^crd/crd^* mutants, suggesting *tnfrsfa* is required for stemness maintenance in neuromasts (Fig. 2G).

Sox2 is has been shown to be expressed in mantle cells (Hernández et al., 2007), and has been hypothesized to be a key factor in maintaining the stem cell niche for the lateral line (Ghysen and Dambly-Chaudière, 2007). Because of the observation that Sox2 positive cells appeared to be reduced in the periphery of the neuromasts (Fig. 2C), we used the Sox2 antibody to test whether mantle cells, marked by *ET20:EGFP*, expressed Sox2 in the lateral line of *tnfrsfa^crd/crd^* mutants. In wild-type embryos, cells double positive for GFP and Sox2 were evident from 48 hpf to 96 hpf. In *tnfrsfa^crd/crd^*mutant embryos, some GFP^+^/Sox2^-^ cells were apparent up to 60 hpf, but after 72 hpf all GFP^+^ cells were Sox2^-^ (Fig. 2H). Therefore, Tnfrsfa activity is necessary to maintain Sox2 activity in the mantle cells but is not necessary for initial activation of *sox2* during development or for maintaining Sox2 activity in the support cells.

We expressed the extracellular domain of zebrafish Tnfrsfa in vitro and produced a polyclonal antibody against Tnfrsfa. Immunofluorescence analysis revealed that Tnfrsfa was not co-localized with the EGFP of *ET20:EGFP*, but enriched in the dorsal compartment of the neuromast, indicating Tnfrsfa functions in support cells but is necessary for maintaining the mantle cells, presumably through maintaining Sox2 activity (Supplementary Fig. S4). The observed data suggests Tnfrsfa functions on the mantle cells in a cell-nonautonomous manner.

### tnfrsfa is necessary for mantle cell proliferation during homeostasis and regeneration

We used BrdU and Sox2 double positive cells to evaluate amplifying cell divisions and BrdU/HCS-1 double positive cell to evaluate differentiating cell divisions during 2 dpf to 3 dpf when the neuromast is growing. The amplifying cell divisions occurred at the peripheral compartment of the neuromast, and the distribution was uniform around both axes (Fig. 3A-B). Compared to wild-types, the amplifying cell divisions occurred less frequently in *tnfrsfa^crd/crd^* mutant neuromasts, whereas the differentiating cell divisions increased significantly in *tnfrsfa^crd/crd^* mutant neuromasts (Fig. 3A-B, 3E). This pattern suggests a premature differentiation of support cells from mantle cells without renewal, which would explain why we detected an increase in hair cells and a decrease of accessory cell in *tnfrsfa* mutants at 2.5 dpf. From 4 dpf to 5 dpf, when the neuromast is in a slower growth state, amplifying cell divisions occur more frequently in the D/V support cells in wild-type neuromasts, consistent with a previous report (Romero-Carvajal et al., 2015). In contrast, *tnfrsfa^crd/crd^*mutant neuromasts exhibited significantly less amplifying and differentiating cell divisions compared to wild-types, which is consistent with the observed decrease in both hair cells and accessory cells at 5 dpf (Fig. 3C-E).

**Figure 3.**
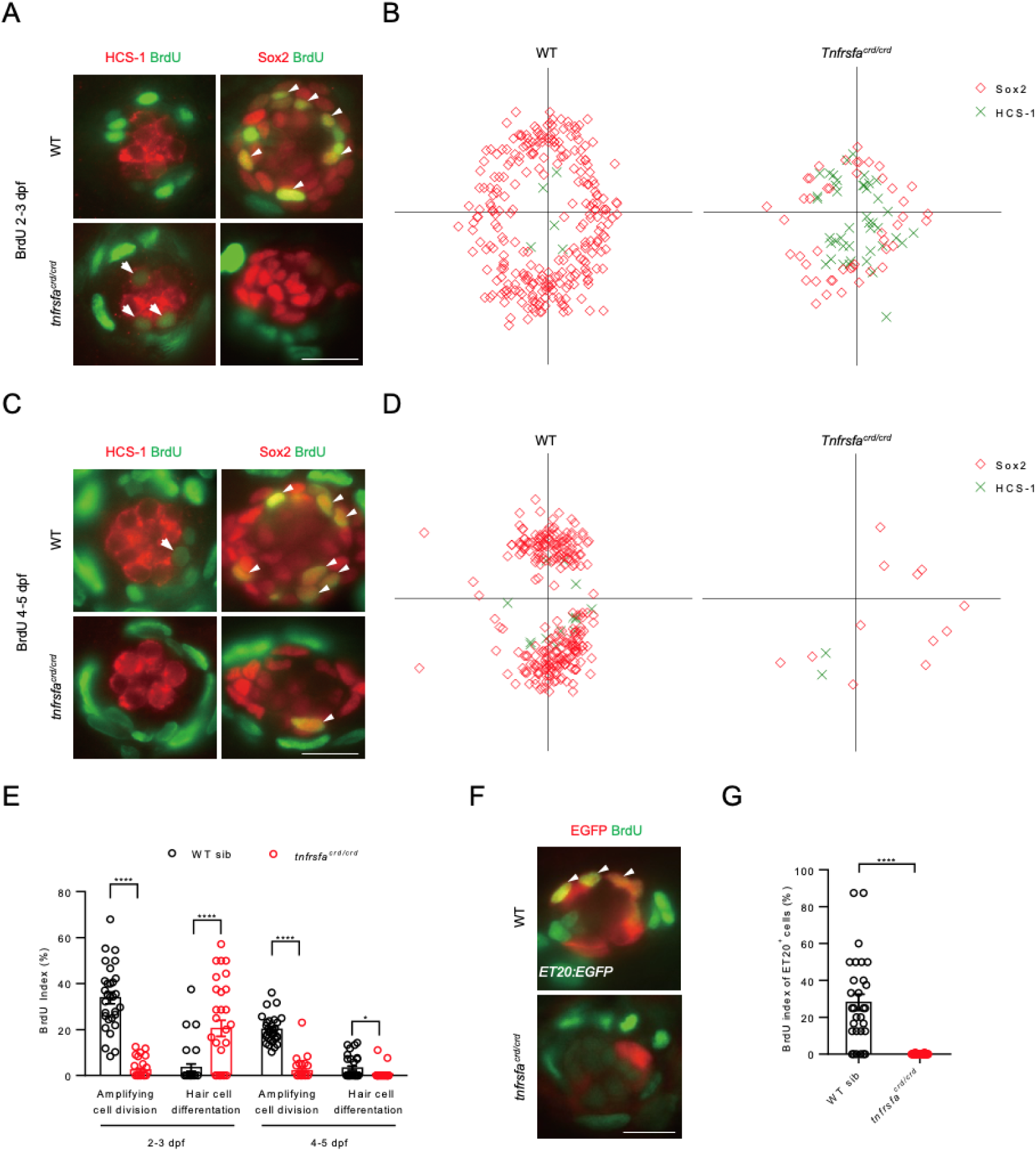
*tnfrsfa* is Essential for Mantle Cell Proliferation During Homeostasis and Regeneration. (A) 24-hour BrdU incorporation in neuromasts from 2 dpf to 3 dpf in wild-type and *tnfrsfa* mutant larvae. Arrowheads indicate BrdU+ support cells; arrows indicate BrdU+ hair cells. Scale bar = 10 μm. (B) Rose diagrams showing the spatial distribution of BrdU+ support cells (red squares) and BrdU+ hair cells (green crosses) in neuromasts. N=30. (C) 24-hour BrdU incorporation from 4 dpf to 5 dpf in wild-type and *tnfrsfa* mutants. Scale bar = 10 μm. (D) Rose diagrams showing BrdU+ cells in neuromasts. N=30. (E) Quantification of BrdU+ support cells and hair cells in wild-type and *tnfrsfa* mutants at different stages. N=30; *, p < 0.05; ****, p < 0.0001. (F) 12-hour BrdU incorporation in neuromasts from *Tg(ET20:EGFP)* following neomycin treatment and LY411575 incubation. Arrowheads mark BrdU+ mantle cells. (G) Quantification of BrdU+ mantle cells during regeneration in wild-type and *tnfrsfa* mutants. N=30; ****, p < 0.0001.

As previously explained, it is possible to reduce the number of support cells during injury by driving them towards hair cell differentiation using the Notch signaling inhibitor LY411575 (Ye et al., 2020). In wild-type neuromasts, if you drive hair cell differentiation with LY411575 and then ablate the hair cells with neomycin, the mantle cells increase cell divisions to restore both support cells and hair cells to their proper numbers. We preformed this “stressed” ablation of hair cells and support cells in *tnfrsfa^crd/crd^* mutants and wild-type siblings. We performed a BrdU incorporation assay to investigate the proliferation of mantle cells in *tnfrsfa^crd/crd^* mutants in the *ET20:EGFP* transgenic background compared to wild-types during the first 12 h after neomycin treatment. The results showed that approximately 28% of mantle cells re-entered the cell cycle in wild-type neuromasts. In contrast, few BrdU positive mantle cells were found in *tnfrsfa^crd/crd^* mutants (Fig. 3F-G). Therefore, mutations in *tnfrsfa* impair the proliferation of mantle cells in response to hair cell injury.

The posterior lateral line neuromasts originate from proto-neuromasts (PNM), which are deposited from the posterior lateral line primordium (PLLP) (Ghysen and Dambly-Chaudière, 2007). To label the PNM and PLLP, *tnfrsfa^crd/crd^*mutants were crossed with the *Tg(-8.0cldnb:LY-EGFP)* transgenic line (*cldnb:EGFP* for short). Similar to mantle cells, the proliferation of the cells in proto-neuromasts was reduced in *tnfrsfa^crd/crd^*mutants compared to wild-types, although the proliferation in the migrating primordium was not changed significantly (Supplementary Fig. S5). The data is consistent with a role for Tnfrsfa in the long-term maintenance of neuromast homeostasis but not directly involved in organization or differentiation of the different cell types.

### NF-***κ***B is activated cell nonautonomously by tnfrsfa in neuromasts

Activation of TNF superfamily receptors usually results in the recruitment of adaptor proteins that trigger the activation of NF-κB, a key transcription factor involved in cytokine production and cell survival which are crucial for tissue regeneration (Aggarwal et al., 2012; Kaltschmidt et al., 2021). To explore whether hair cell regeneration is correlated with NF-κB activation, we examined NF-κB activation in response to neuromast injuries using the transgenic zebrafish line *Tg(NFkB:EGFP)* (*NFKB:EGFP* for short). NF-κB signaling was only strongly activated in mantle cells when both support cells and hair cells were ablated using LY411575 and neomycin, rather than with hair cell loss alone in the neomycin treatment (Fig. 4A). This indicates that while there is always some NF-κB activity in mantle cells, strong NF-κB activation is a hallmark of severe neuromast injury when the mantle cells have to enter mitosis in significant numbers.

**Figure 4.**
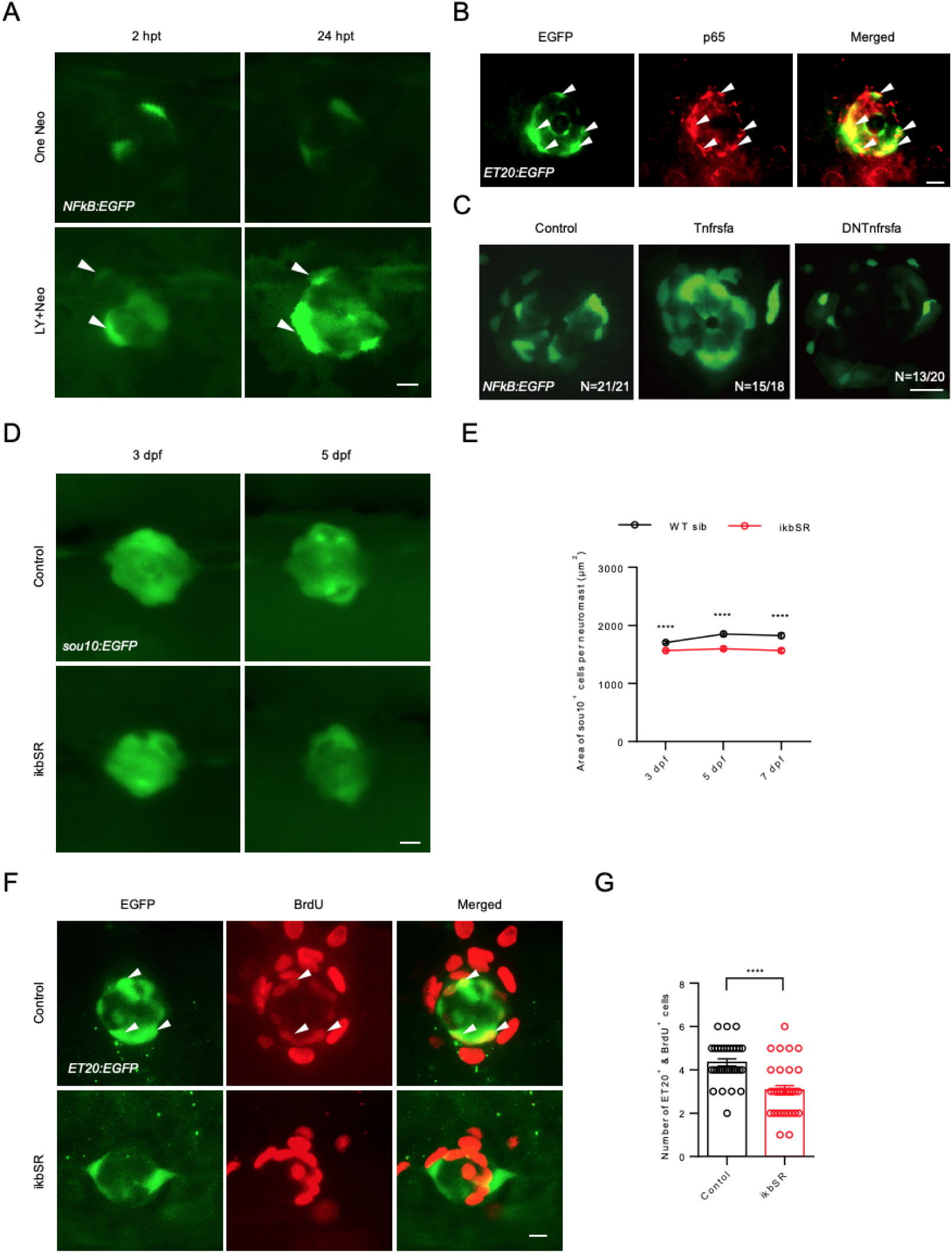
NF-κB Signaling is downstream of *tnfrsfa* in Neuromast Development. (A) Live imaging of neuromasts from *Tg(NFkB:EGFP)* larvae treated with neomycin (One-Neo) or neomycin plus LY411575 (LY and Neo). Arrowheads indicate activated NF-κB signaling in mantle cells. Scale bar = 10 μm. (B) Immunostaining of EGFP (green) and p65 (red) in neuromasts of *Tg(ET20:EGFP)* larvae at 5 dpf. Arrowheads show co-localization of EGFP (green) and p65 (red). Scale bar = 10 μm. (C) Live imaging of neuromasts from *Tg(NFkB:EGFP)* (Control), *Tg(NFkB:EGFP); Tg(hsp70l:Tnfrsfa-mcherry)* (Tnfrsfa) and *Tg(NFkB:EGFP); Tg(hsp70l:DNTnfrsfa-mcherry)* (DNTnfrsfa) following heat-shock. Scale bar = 10 μm. (D) Live imaging of neuromasts from Tg(sou10:EGFP) (Control) and *Tg(hsp70l:ikbSR-mcherry)* (ikbSR) following heat-shock at 3 dpf and 5 dpf. Scale bar = 10 μm. (E) Quantification of neuromast area in *Tg(sou10:EGFP)* lines with (red) or without (black) ikbSR. N=40-60; ****, p < 0.0001. (F) 12-hour BrdU incorporation in *Tg(ET20:EGFP)* neuromasts after LY+Neo treatment in both wild-type and *tnfrsfa* mutants. Arrowheads show co-localization of EGFP and BrdU. Scale bar = 10 μm. (G) Quantification of BrdU^+^ mantle cells after LY+Neo treatment. N=32; ****, p < 0.0001.

We used an NF-κB subunit p65 antibody in combination with the *ET20:EGFP* transgene, to investigate the relationship between NF-κB signaling and mantle cells. In the mature neuromasts at 5 dpf, most p65 was co-localized with EGFP in *ET20:EGFP* transgenic fish (Fig. 4B), which is consistent with the distribution of *NFkB:EGFP* transgenic activity in the neuromast (Kanther et al., 2011; Lush et al., 2019).

To investigate whether NF-κB signaling acts downstream of Tnfrsfa, we performed in situ hybridization to measure the expression of *nfkbib,* a known target of NF-κB signaling (Schreiber et al., 2006). We found high *nfkbib* expression in wild-type neuromasts, but the expression of *nfkbib* was decreased in *tnfrsfa* mutants (Supplementary Fig. S6). While *nfkbib* is a target of NF-κB signaling, it is an inhibitor of NF-κB and it was expressed throughout the support cells, not in the mantle cells. We interpret this result as Tnfrsfa activates NF-κB, but steady Tnfrsfa signaling in the support cells ultimately inhibits NF-κB activity in the support cells through a feedback loop. The support cells must then send a signal back to the mantle cells that maintains NF-κB activity in the mantle cells.

We generated transgenic lines to express wild-type Tnfrsfa or a dominant-negative version of Tnfrsfa (lacking the Death Domain) under the transcriptional control of the *heat-shock protein 70l* (*hsp70l*) promoter. Heat-shocking *Tg(hsp70l:Tnfrsfa-mCherry)* transgenic larvae (*hsp:Tnfrsfa* for short) led to a significant increase of NF-κB signaling activity in mantle cells shown by the *Tg(NFkB:EGFP)* transgene, while heat-shocking *Tg(hsp:DNtnfrsfa-mcherry)* transgenic larvae (*hsp:DNTnfrsfa* for short) resulted in significantly reduced NF-κB signaling activity in mantle cells (Fig. 4C). This indicates that NF-κB signaling in the mantle cells is induced by Tnfrsfa expression in neuromasts. It is notable that widespread expression of Tnfrsfa was not sufficient for activation of NF-κB activity broadly since the increase in activity was still limited to mantle cells despite the receptor being ubiquitously expressed.

To establish the relationship between NF-κB signaling and hair cell regeneration, we generated a *Tg(hsp70l:ikBSR-mCherry)* transgene (*hsp:ikBSR* for short) expressing a super-repressor of NF-κB upon heat-shock. The effectiveness of ikBSR was confirmed by reduced NF-κB activity in the *NFkB:EGFP* reporter line following two heat-shocks (Supplementary Fig. S7). The overexpression of ikBSR led to a significant reduction in hair cells shown *by Tg(Brn3c:EGFP)* (*Brn3c:EGFP* for short) (Supplementary Fig. S8) and support cells shown by *sou10:EGFP* (Fig. 4D-E) in neuromasts at 5 dpf. Notably, mantle cell proliferation, assessed by a BrdU assay using the *ET20:EGFP* reporter, was significantly reduced during hair cell regeneration after support cell and hair cell injury (LY followed by neomycin treatment) (Fig. 4F-G). Thus, down-regulation of NF-κB signaling mimics the *tnfrsfa* mutant phenotype, with impaired mantle cell proliferation. The results indicate that NF-κB signaling mediates the role of Tnfrsfa in neuromasts. Because Tnfrsfa is primarily expressed in dorsal support cells, but NF-κB activity is detected in the mantle cells, Tnfrsfa must be activating NF-κB in the mantle cells in a cell nonautonomous manner.

### tnfsf10 and tnfsf10l3 are required for neuromast development in zebrafish

*tnfsf10* (also known as *zDLa*) and *tnfsf10l3* (also known as *zDL1b*) have been previously reported to be highly expressed in developing neuromasts. Both Tnfsf10 orthologs have been shown to act as a ligand for Tnfrsfa (Eimon et al., 2006). Published single cell data confirms both are expressed in the neuromast, with *tnfsf10* strongly enriched in the mantle cells and *tnfsf10l3* expressed at a lower level and more broadly distributed across all the cell types of the neuromast (Baek et al., 2022).

To further investigate their roles in neuromast development, we targeted the coding sequences of the TNF homology domain in both *tnfsf10* and *tnfsf10l3* using CRISPR/Cas9 and generated mutant lines for each (Supplementary Fig. S9; Fig. 5A). Phenotypic analyses revealed reductions in both hair cells and support cells in *tnfsf10* and *tnfsf10l3* mutants compared to wild-type controls (Fig. 5B-D).

**Figure 5.**
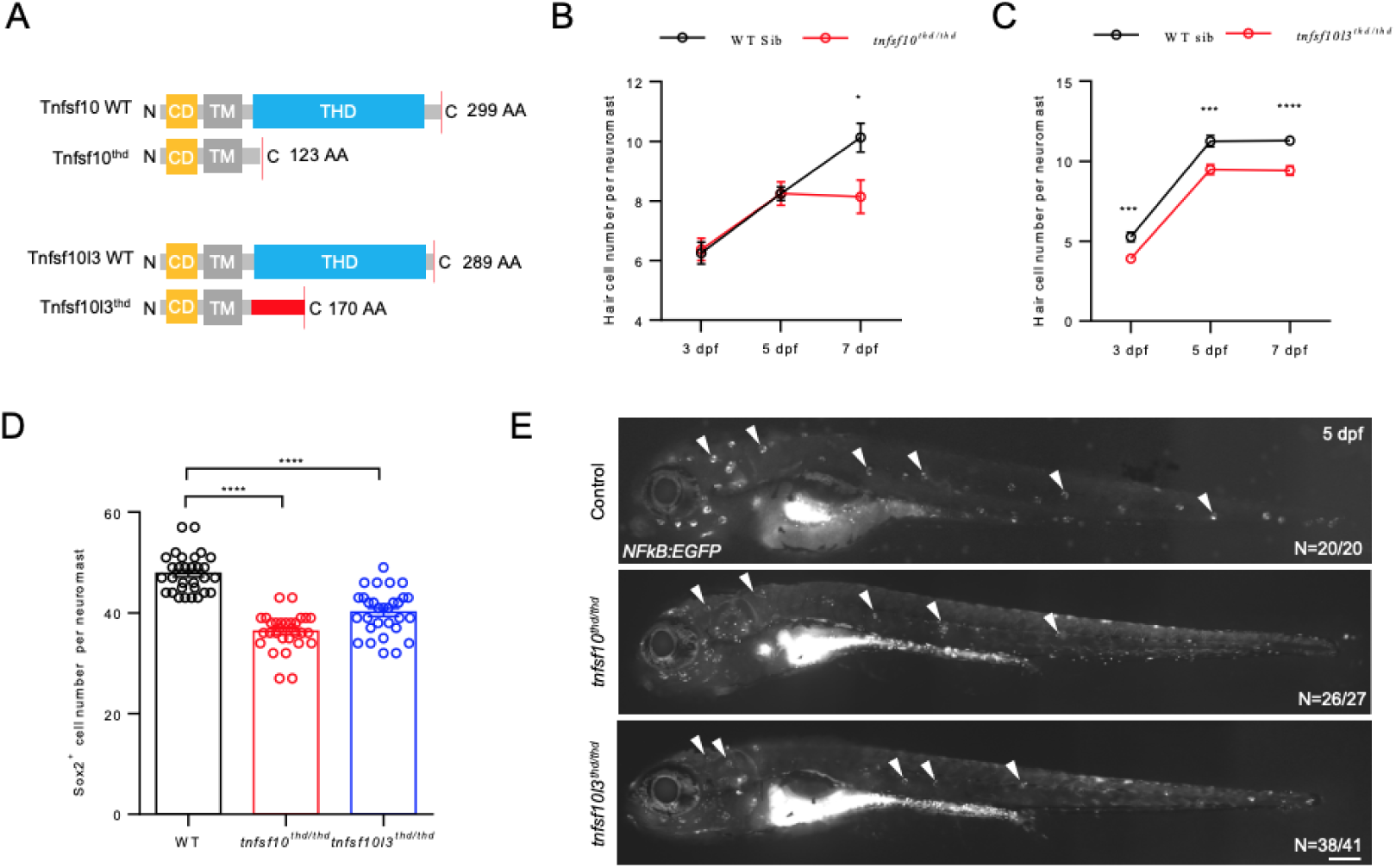
*tnfsf10* and *tnfsf10l3* are Required for Neuromast Development in Zebrafish. (A) Schematic showing wild-type and mutant *Tnfsf10* and *Tnfsf10l3* generated via CRISPR/Cas9. CD, cytoplasmic domain; TM, transmembrane domain; THD, TNF homology domain. Red box indicates missense sequence. (B) Quantification of hair cells in posterior lateral line neuromasts of *tnfsf10* mutants and wild-type siblings using Yo-Pro-1 staining. N=10. (C) Quantification of hair cells in *tnfsf10l3* mutants using Yo-Pro-1 staining. N=6-10. (D) Quantification of accessory cells in neuromasts of *tnfsf10* and *tnfsf10l3* mutants using Sox2 immunostaining at 5 dpf. N=30. (E) Live imaging of *Tg(NFkB:EGFP)* larvae at 5 dpf in wild-type, *tnfsf10^thd/thd^*, and *tnfsf10l3^thd/thd^* mutants. Arrowheads indicate neuromasts. Scale bar = 200 μm.

Furthermore, we observed that the deficiency of either *tnfsf10* or *tnfsf10l3* led to a marked reduction in NF-κB signaling activity as demonstrated by *NFkB:EGFP* reporter line (Fig. 5E). These results indicate that Tnfsf10 and Tnfsf10l3 are both involved in the lateral line development and hair cell maintenance through Tnfrsfa and NF-κB signaling. The phenotype of *tnfsf10* more closely matches the *tnfrsfa* phenotype (i.e. development appears normal at first but hair cell number levels off prematurely) and is more strongly and specifically expressed in the mantle cells (Baek et al., 2022) making its interactions with *tnfrsfa* likely to be the driving interaction in the failure to maintain the mantle cells in a stem cell fate.

Using the single cell RNA-seq neuromast data of Baek *et al*., we identified other members of TNF receptor superfamily that were also expressed in zebrafish neuromast cells. So, we evaluated the role of *tnfrsf19*, *tnfrsf21*, and *hdr* by generating F0 mosaic mutant zebrafish (“crispants”). The results showed that the knockouts of *tnfrsf19*, *tnfrsf21*, and *hdr* did not affect the NF-κB signaling in the neuromast, although hair cell number was decreased in the *hdr-crispants* (Supplementary Fig. S10).

### tnfrsfa regulates neuromast development independent of Notch signaling

Notch signaling is known to regulate support cell identity within neuromasts and influence hair cell regeneration through lateral inhibition (Romero-Carvajal et al., 2015; Thomas and Raible, 2019). To assess the relationship between Tnfrsfa and Notch signaling, we performed *in situ* hybridization to show the expression of Notch signaling target genes in the mutant background. The results showed that *her4.1* and *notch3* in *tnfrsfa^crd/crd^*larvae were comparable to wild-type levels (Supplementary Fig. S11). Furthermore, we analyzed the activity of the Notch signaling reporter line, *Tg(tp1:dGFP)* (*tp1:dGFP* for short), in response to changes of Tnfrsfa signaling. Activation or inhibition of Tnfrsfa signaling with *hsp:Tnfrsfa* and *hsp:DNtnfrsfa*, respectively did not significantly alter the activity of *tp1:dGFP* which specifically labels hair cells (Fig. 6A).

**Figure 6.**
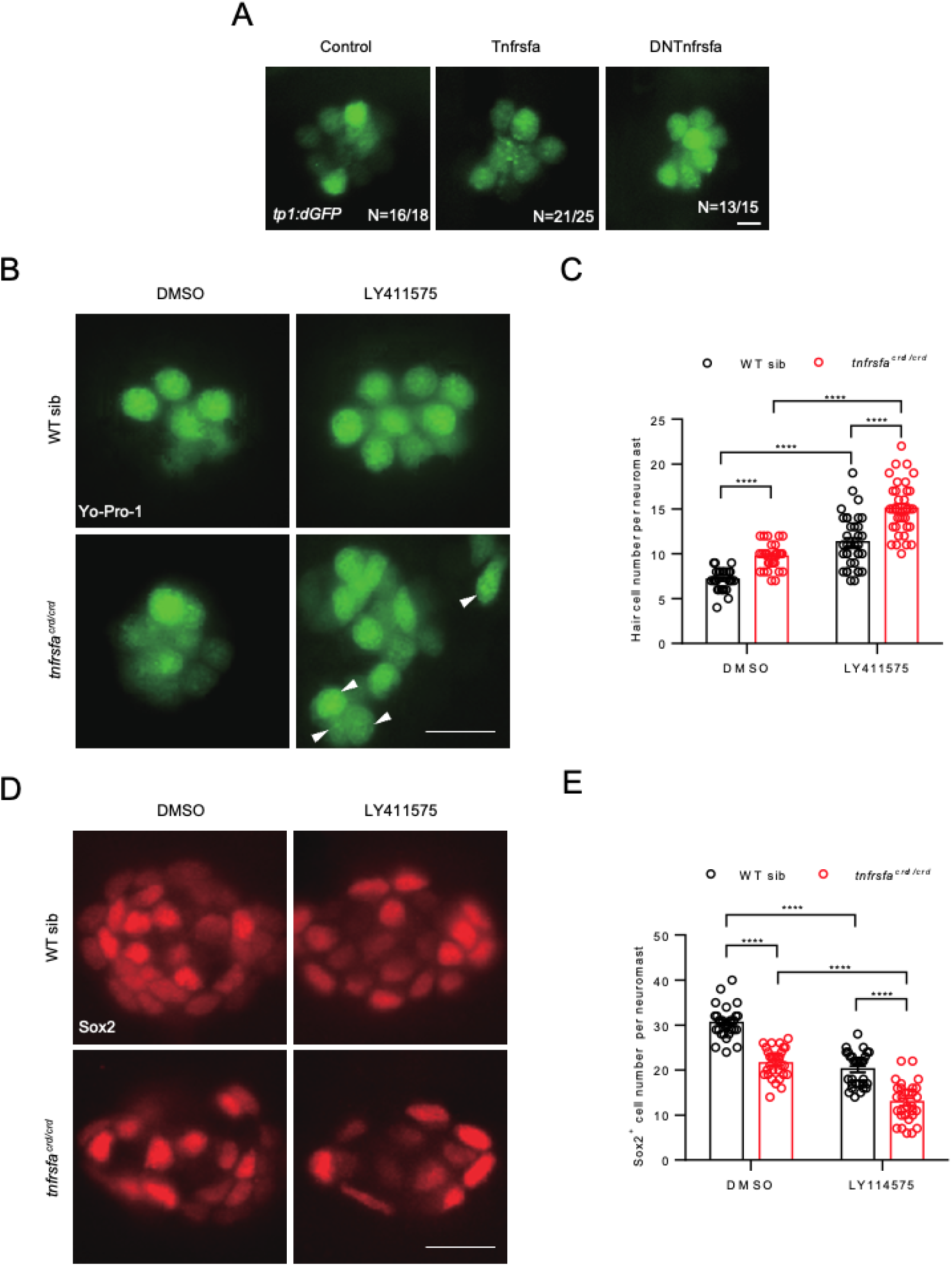
Tnfrsfa Regulates Neuromast Development Independently of Notch Signaling. (A) Live imaging of neuromasts at 5 dpf from *Tg*(*tp1:dGFP*) which labels hair cells (Control), *Tg*(*tp1:dGFP*)*; Tg*(*hsp70l:Tnfrsfa-mcherry*) (Tnfrsfa) and *Tg*(*tp1:dGFP*)*; Tg*(*hsp70l:DNTnfrsfa-mcherry*) (dominant negative Tnfrsfa) following heat-shock. No differences were detected. Scale bar = 10 μm. (B) Live imaging of neuromasts at 3 dpf visualized by Yo-Pro-1 staining in wild-type and *tnfrsfa* mutant larvae treated with DMSO or LY411575 for 2 days. Arrowheads indicate peripheral hair cells. Both mutant and wild-type siblings increased hair cell number in the presence of Notch inhibitors to the same degree. Scale bar = 10 μm. (C) Quantification of hair cells in each neuromast of wild-type and *tnfrsfa* mutant larvae treated with DMSO or LY411575 for 2 days. N = 36; ****, p < 0.0001. (D) Immunostaining of neuromasts with Sox2 antibody at 3 dpf in wild-type and *tnfrsfa* mutant larvae treated with DMSO or LY411575 for 2 days. (E) Quantification of support cells in each neuromast of wild-type and *tnfrsfa* mutant larvae treated with DMSO or LY411575 for 2 days. N = 30; ****, p < 0.0001.

Both Notch signaling inhibition and *tnfrsfa* mutations result in precocious hair cells at the expense of accessory cell decrease during neuromast development. We examined the effects of simultaneous inhibition of Notch and Tnfrsfa on the growth of hair cells and support cells. Wild-types and *tnfrsfa* mutants were treated with LY411575 from 1 dpf to 3 dpf. In LY411575-treated *tnfrsfa* mutants, the number of hair cells was higher than untreated *tnfrsfa* mutants and LY411575-treated wild-types, with no significant interaction between Tnfrsfa deficiency and Notch suppression on hair cell induction (P = 0.65) (Fig. 6B-C) suggesting an additive effect between the two pathways. Additionally, support cell numbers were increased in LY411575-treated *tnfrsfa* mutants compared to untreated *tnfrsfa* mutants and LY411575-treated wild-types, with no significant interaction observed between Tnfrsfa deficiency and Notch suppression (P = 0.33) (Fig. 6D-E). These findings suggest that Tnfrsfa and Notch signaling play distinct roles in maintaining neuromast homeostasis. A simple explanation would be that Tnfrsfa signaling is involved in preserving the ability of mantle cells to amplify and replace support cells, while Notch signaling is involved in determining cell fates within the support cells and hair cells, that would place Tnfrsfa upstream of Notch signaling.

## Discussion

Using various mutant and transgenic zebrafish lines, we showed that the inflammatory signaling pathway involving Tnfsf10, Tnfsf10l3, Tnfrsfa, NF-κB and Sox2 is crucial for maintaining the stem cell niche of lateral line neuromasts allowing for the juvenile and adult expansion of neuromast numbers and to maintain the capacity for hair cell regeneration within the neuromasts. This pathway specifically maintains the ability of mantle cells to proliferate during development and regeneration which we believe represents the bona fide stem cell niche for the lateral line neuromasts.

Both hair cells and entire neuromasts can be continually replaced in the fish lateral line throughout their lives demonstrating the existence of stem cells within the neuromast (Behra et al., 2009; Cruz et al., 2015; Dufourcq et al., 2006; Ledent, 2002; Pinto-Teixeira et al., 2015; Song et al., 1995; Wada et al., 2013). We found that the loss of *tnfrsfa* results in the complete disappearance of lateral line sensory organs in adult zebrafish and a significant impairment in hair cell regeneration following severe or repeated injury.

The analysis of accessary cells revealed that peripheral mantle cells in mutant neuromasts are severely reduced or absent in *tnfrsfa* mutants as demonstrated by *tnfrsfa* mutants lacking *ET20:EGFP* positive cells. The number, morphology, and function of support cells remained unchanged at early larval stages, indicating that the reduction in support cells is a long-term consequence of mantle cell loss and the Tnfrsfa signaling is not essential for support cell or hair cell differentiation or function.

Previous reviews have proposed that mantle cells serve as stem cells in neuromast (Denans et al., 2019; Ghysen and Dambly-Chaudière, 2007). Our study provides direct evidence that peripheral cells, particularly mantle cells, act as the stem cells of the neuromast. Consistent with our findings, mantle cells in both zebrafish and medaka have been shown to differentiate into all cell types within the neuromast (Seleit et al., 2017). Recent evidence in medaka suggests post-larval addition of neuromasts occurs by mantle cells delaminating from existing neuromasts, migrating to a new location, and forming new neuromasts (Gross et al., 2022). This is consistent with our claim that the Sox2^+^ mantle cells are the stem cell niche in zebrafish and with our observations that early neuromast development occurs relatively normally in *tnfrsfa* mutants, but further expansion of the lateral line network does not occur, presumably because the loss of stem cell identity in the mantle cells would prevent the delamination and expansion of the lateral line organ in adult fish. Moreover, mantle cells remain quiescent under normal physiological conditions and are activated to proliferate and differentiate into support cells only after severe injury (Romero-Carvajal et al., 2015; Thomas and Raible, 2019). This behavior parallels that of long-term stem cells observed in other tissues, such as the intestine, hematopoietic systems, and neural tissues (Barker et al., 2008; Kobayashi et al., 2019; Pinho and Frenette, 2019).

As others have proposed (Romero-Carvajal et al., 2015) we believe the central support cells act as immediate precursors to hair cells, while D/V support cells serve as transit-amplifying cells responsible for replenishing support cell numbers. Mantle cells represent the *bona fide* neuromast stem cells that self-renew and replenish the D/V support cells (Fig. 7). As previously reported, A/P support cells also remain quiescent during homeostasis and modest hair cell injury (Thomas and Raible, 2019). Specific ablation of mantle cells and A/P support cells may further elucidate the roles of these cells during homeostasis and regeneration within the neuromast.

**Figure 7.**
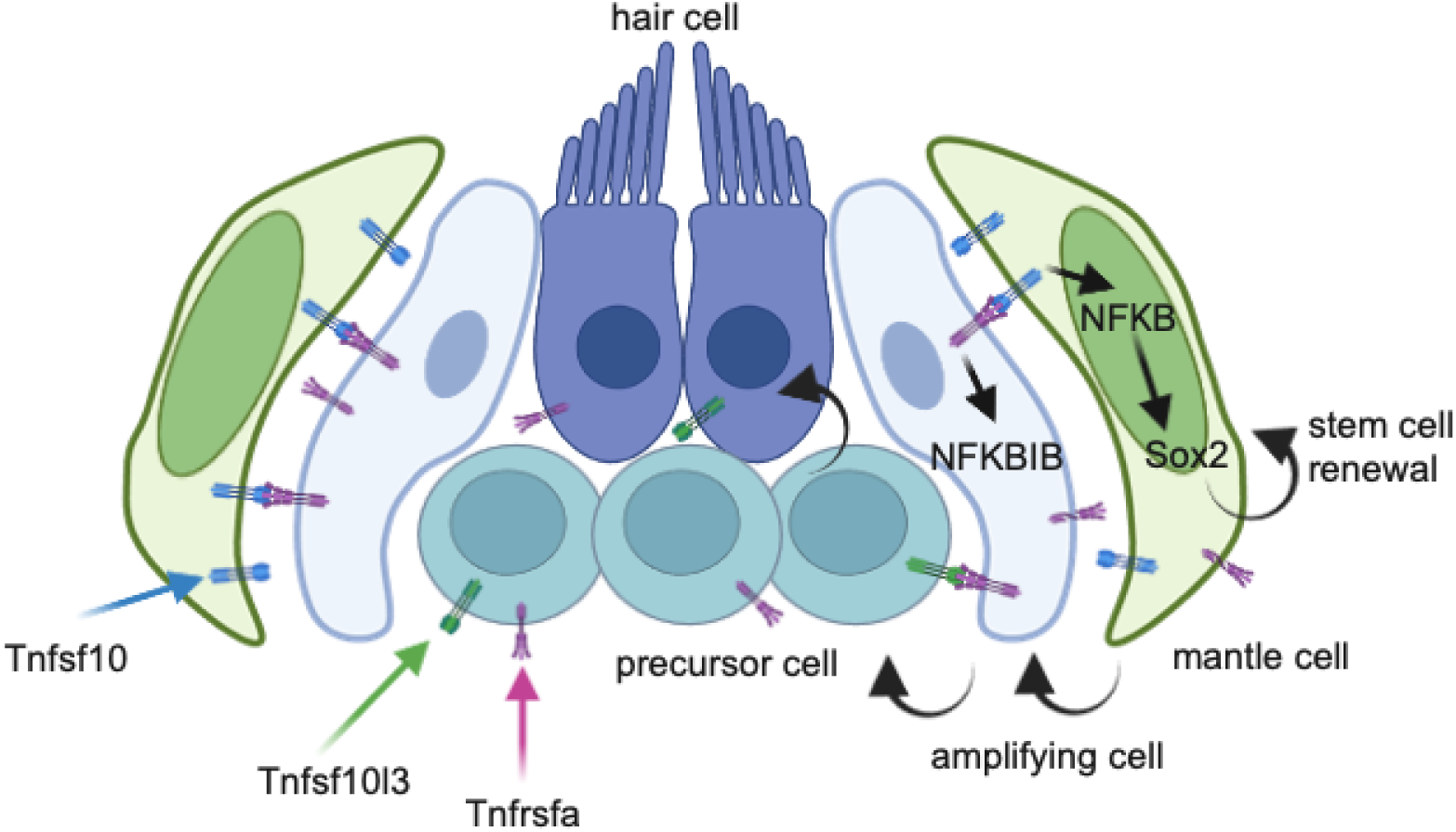
Schematic diagram that illustrates a model for how the Tnfrsfa-NF-κB-Sox2 pathway maintains mantle cell self-renewal in neuromasts. The mantle cells express Tnfsf10 at high levels and support cells throughout the neuromast but concentrated in the dorsal area express Tnfrsfa. Tnfsf10 activates NFKBIB activity in the support cells and reverse signals from the transmembrane version of Tnfsf10 activate NFKB and Sox2 in the mantle cells. Sox2 expression allows for mantle cells to maintain stem cell identity and self-renew without premature differentiation into support cells or hair cells. After injury, mantle cells can become support cells to replenish the pool of transit-amplifying cells that can become hair cells when needed.

The conserved inflammatory signaling pathways, such as MAPK/AP-1, Hippo/YAP, Jak/Stat, and NF-κB, have critical roles in tissue regeneration both through inflammation directly or local use of the same signaling pathways by other cells (Eming et al., 2017; Favier and Nikovics, 2023; Karin and Clevers, 2016). Previous studies have highlighted the involvement of inflammation regulators like Stat3, HSP60, JNK, and AP-1 in hair cell regeneration in the zebrafish neuromast and inner ear (Baek et al., 2022; He et al., 2016; Liang et al., 2012; Pei et al., 2018). Our research showed that mutations in *tnfrsfa* or the overexpression of ikbSR, a known repressor of the NF-κB signaling pathway, significantly impaired hair cell regeneration following severe neuromast damage. This finding parallels the role of the TNFα/TNFR1 pathway in fin and retinal regeneration (Nelson et al., 2013; Nguyen-Chi et al., 2017). There are also interesting parallels between our findings and those of Espín-Palazón et al., they showed that definitive hematopoietic stem cells in zebrafish were induced from the arterial endothelium through Tnfa/Tnfr2 signaling (Espín-Palazón et al., 2014). Similar to our results, the induction is done through NF-κB and is independent of the Notch pathway which is also required in parallel for establishing hematopoietic stem cells.

Inflammation supports tissue regeneration by managing infections, clearing damaged cells, and creating an appropriate niche for stem cells (Eming et al., 2017; Favier and Nikovics, 2023; Karin and Clevers, 2016). During hair cell regeneration, Stat3 and JNK influence the proliferation and differentiation of support cells, which act as transit amplifying cells within the neuromast (He et al., 2016; Jiang et al., 2014; Liang et al., 2012). Our data indicate that NF-κB signaling in mantle cells is only increased noticeably by significant support cell depletion, and either loss of *tnfrsfa* or ikbSR overexpression decreased mantle cell proliferation. Thus, Tnfrsfa-NF-κB signaling is critical for maintaining and proliferating the stem cells of the neuromast.

Tnfsf10, Tnfsf10l3, and Tnfrsfa in zebrafish are homologous to TRAIL and its receptor in mammals (Eimon et al., 2006). Although death receptors are present in the mammalian cochlea, TRAIL does not effectively promote hair cell regeneration there, but leads to hair cell death and neuronal degeneration instead (Hayashi et al., 2021; Kao et al., 2016). We did not detect any disruptions of hair cell development in the adult zebrafish inner ear (which can also regenerate) in the absence of Tnfrsfa activity so the establishment of stem cells for the lateral line may arise from the distinct structure and nature of the cells of the fish neuromast versus how the niche is established in the inner ear.

In contrast to the later neuromast deficiencies observed in *tnfrsfa* mutant adults, *tnfrsfa* mutants exhibited an increased number of hair cells at early larval stages compared to wild-type, indicating premature differentiation of hair cells. This phenotype resembles that of *mindbomb* mutant zebrafish, which also display an excess of hair cells in the inner ear and neuromasts (Haddon et al., 1998; Itoh and Chitnis, 2001). It has been reported that Notch signaling plays a crucial role in maintaining support cells in a quiescent state and suppressing their differentiation into hair cells but does not affect the proliferation and differentiation of mantle cells (Romero-Carvajal et al., 2015; Thomas and Raible, 2019). Our findings indicate that *tnfrsfa* mutants treated with a γ-secretase inhibitor produced more hair cells compared to both untreated *tnfrsfa* mutants and γ-secretase inhibitor-treated wild-types. This suggests that the Notch and NF-κB signaling pathways have parallel but distinct roles in neuromast development and homeostasis.

Tnf superfamily signaling plays a role in hematopoietic and neural stem cell functions (Corsini et al., 2009; Espín-Palazón et al., 2014). Additionally, dysregulation of *TNFRSF10A* is linked to age-related macular degeneration in humans (Arakawa et al., 2011; Mori et al., 2022). The interactions between mantle and support cells in neuromasts resembles the relationship between intestinal stem cells and Paneth cells, which support the stem cells through niche signals (Karin and Clevers, 2016). Tnfsf10 and Tnfrsfa have been shown to inhibit muscle cell differentiation (Kim et al., 2020) suggesting the complex could be involved in muscle stem cell maintenance. Our model is that Tnfrsfa-positive support cells activate the highly positive Tnfsf10 mantle cells and activate intracellular signals that regulate mantle cell proliferation via induction of NF-κB signaling in the mantle cells through a mechanism similar to Delta-Notch signaling (Fig. 7). Tnf super family members possess transmembrane domains and have been shown to participate in “reverse signaling,” i.e. when the membrane-bound ligand interacts with the receptor, the ligand also sends an intracellular signal inducing changes in the cell expressing the ligand (Lee et al., 2019). This reverse signaling has been demonstrated for Tnfsf10 (Chou et al., 2001) including induction of NF-κB (Huang et al., 2011). We showed this NF-κB activity is necessary for *sox2* expression in the mantle cells which maintains the stem cell identity. In some cancers, NF-κB activity is linked to Sox2 expression and maintenance of cancer stem cells (Vazquez-Santillan et al., 2016; Zakaria et al., 2018). Consistent with this hypothesis is that when we induce Tnfrsfa broadly with a heat shock, we still only see increased NF-κB activity in Tnfsf10 positive mantle cells. Reverse signaling is the parsimonious explanation of how Tnfrsfa activity in the support cells induces NF-κB and Sox2 in the mantle cells, but it needs to be further explored.

## Materials and Methods

### Zebrafish husbandry and maintenance

The zebrafish lines used in this work include the wild-type line of AB, and several transgenic lines of *Et(krt4:EGFP)sqet20* (*ET20:EGFP* for short) (Parinov et al., 2004), *Tg(tnks1bp1:EGFP)* (*SCM1:EGFP* for short) (Behra et al., 2012), *Tg(-8.0cldnb:LY-EGFP)* (*cldnb:EGFP* for short) (Haas and Gilmour, 2006), *Et(gata2a:EGFP)sou10* (*sou10:EGFP* for short) (Fan et al., 2022), *Tg(Brn3c:GAP43-GFP)s356t* (*Brn3c:EGFP* for short) (Xiao et al., 2005), and *Tg(tp1-MmHbb:EGFP)um14* (*tp1:EGFP* for short) (Clark et al., 2012). Adult zebrafish were maintained in a circulating water system at 28°C with a 14-hour light/10-hour dark cycle. Embryos were produced by natural mating and staged according to previous work (Kimmel et al., 1995). All animal procedures were performed under the regulations of the Animal Ethics Committee of Shanghai Ocean University.

### Transgenic line generation

The hsp70l:Tnfrsfa-mCherry and hsp70l:DNtnfrsfa-mcherry constructs were generated via the Gateway Tol2 system (Thermo Fisher). pME-Tnfrsfa and pME-DNTnfrsfa (the whole Tnfrsfa coding sequence and the Tnfrsfa lacking the DD were amplified and cloned from zebrafish embryonic cDNA) constructs were generated via BP recombination, and then the destination constructs were generated via LR recombination of p5E-hsp70l, pME-Tnfrsfa, pME-DNTnfrsfa, p3E-mCherrypA, and pDestTol2pA vectors (Kwan et al., 2007). The phsp70l:ikbSR-mcherry construct was generated with the ClonExpress II One Step Cloning Kit (Vazyme, Nanjing China). The ikbSR fragments with mutations in S35A and S39A of iκB were cloned from zebrafish embryonic cDNA with primers containing substituted bases. The Tol2 plasmid backbone sequence containing the *hsp70l* promoter and mCherry was amplified from the phsp70l:Tnfrsfa-mCherry construct. The pET28a-CRD construct was generated by traditional cloning methods, by cloning the extracellular domain coding sequence of Tnfrsfa from zebrafish embryonic cDNA into pET28a plasmid digested with *Nco I* and *Xho I.* All plasmids were extracted using maxi prep (Tiangen, China) prior to injection.

To produce the *Tg(NFkB:EGFP)*, *Tg(hsp70l:Tnfrsfa-mCherry)*, *Tg(hsp70l:DNtnfrsfa-mcherry)* and *Tg(hsp70l:ikbSR-mcherry)* transgenic lines, recombined plasmids containing Tol2 ITRs were injected into one-cell stage embryos derived from the AB strain along with transposase mRNA. Founders with GFP or mCherry positive cells were identified by crossing with wild-type fish. The positive carriers have been maintained for over three generations.

### Gene knockout using CRISPR/Cas9

Zebrafish mutants were generated using CRISPR/Cas9 as previously described (Duan et al., 2024; Varshney et al., 2016). Single guide RNA (sgRNA) targets for *tnfrsfa*, *tnfsf10*, and *tnfsf10l3* were designed using the CRISPRScan track in the UCSC genome browser (Moreno-Mateos et al., 2015). Target-specific oligos are listed in Supplementary Table 1. sgRNA transcription templates were prepared by PCR and subsequently *in vitro* transcribed using the T7 High Yield RNA Synthesis Kit (Vazyme, Nanjing, China). Cas9 mRNA was *in vitro* transcribed with mMessage mMachine T3 Transcription Kit (Thermo Fisher, USA) from pT3TS-nCas9n (addgene # 46757) linearized by *Xba I*. Each wild-type embryo was injected with a mixture of approximately 300 pg of Cas9 mRNA and 50 pg of gRNA at the one-cell stage. Indel efficiency was evaluated using the CRISPR-STAT method (Carrington et al., 2015). The progeny of founder fish with desired mutations were raised to adulthood, genotyped by fluorescent PCR followed by capillary electrophoresis, and sequenced to determine the predicted effect on the encoded protein.

### Whole-mount in situ hybridization

Whole-mount in situ hybridization was performed as described by Thisse and Thisse (Thisse and Thisse, 2008). Antisense RNA probes were synthesized using DIG RNA Labeling Mix (Roche, USA) and T7 High Yield RNA Transcription Kit (Vazyme, Nanjing, China). The primers used to clone the transcription template for probes are listed in Supplementary Table 1. Hybridization was performed using 1-10 ng/μl probe at 65°C for at least 16 h. Anti-DIG-AP antibody (Roche, Swiss) diluted to 1:5000 was used for detecting the DIG labeled probe. Samples were stained with BM Purple (Roche, Swiss), immersed in glycerol, and photographed with a Nikon Eclipse 80i microscope.

### Polyclonal antibody preparation of Tnfrsfa

Expression of the extracellular domain of zebrafish Tnfrsfa (AA1-AA160) with a His-tag was induced by adding isopropyl β-D-thiogalactoside (IPTG) to the culture of *Escherichia coli* BL21 transformed with pET28a-CRD. The recombinant protein was purified with a Ni-NTA resin column (Sangon, China) and verified by SDS-polyacrylamide gel electrophoresis (PAGE). A rabbit polyclonal Tnfrsfa antibody was generated against the recombinant protein.

### Immunohistochemistry

Immunohistochemistry was performed as described previously (Zhang et al., 2024). Zebrafish larvae were fixed with 4% PFA and washed with PBDT (PBS with 1% DMSO and 0.1% Tween-20). The primary antibodies include rabbit anti-Sox2 (1:200 dilution, Sigma), rabbit anti-HCS-1 antibodies (1:200 dilution, ABclonal), rabbit anti-p65 (1:200 dilution, WanleiBio), mouse anti-BrdU with Alexa Fluor 546 conjugate (1:200 dilution, Invitrogen), and mouse anti-EGFP antibody (1:500, Invitrogen). The secondary antibodies include goat anti-rabbit Alexa Fluor 594 conjugated IgG (1:200, Invitrogen, USA) and goat anti-mouse Alexa Fluor 488 conjugated IgG. After washing with PBDT, images were acquired under a Zeiss Axio Observer Z1 fluorescent microscope using an AxioCam camera.

### BrdU and EdU assays

Cell proliferation in neuromasts was assessed using Bromodeoxyuridine (BrdU, Sigma-Aldrich, USA). Larvae were incubated in 10 mM BrdU with 1% DMSO in Holtfreter’s buffer, then fixed overnight in 4% paraformaldehyde (PFA) at 4°C. BrdU was detected with mouse anti-BrdU antibody (1:200 dilution; Santa Cruz, USA). After five washes with PBDT, the samples were incubated with anti-mouse Alexa Fluor 488 conjugated IgG (1:200 dilution; Invitrogen, USA).

Click-iT EdU kit (Invitrogen, USA) was used to examine the cell proliferation in the posterior lateral line primordium. Zebrafish larvae were collected at 30 hpf and incubated in an EdU solution (500 μM EdU, 10% DMSO, 85% Holtfreter’s buffer) at 28°C for 1 h. Larvae were then fixed in 4% PFA overnight, washed and incubated in Click-iT EdU Alexa Fluor 647 according to the manufacturer’s instructions.

### Yo-Pro-1 and alkaline phosphatase

The vital dye YO-PRO-1 Iodide (Molecular Probes, USA) was used to stain hair cells in neuromasts. Larvae were incubated in 2 μM YO-PRO-1 diluted in Holtfreter’s buffer for 1 h in the dark. After washing with Holtfreter’s buffer 3 times, the larvae were anesthetized with 0.1 mg/ml tricane (MS-222, Sigma) and observed under a fluorescent inverted microscope.

For alkaline phosphatase staining, larvae were fixed overnight in 4% PFA at room temperature, then bleached in 10% H_2_O_2_ with 5% formamide. After 3 washes with PBS, larvae were incubated in alkaline phosphatase buffer (50 mM MgCl_2_, 100 mM NaCl, 100 mM Tris pH 9.5 and 0.1% Tween-20) for 15 min, followed by NBT/BCIP staining solution (Roche, USA) at room temperature in the dark. The staining reaction was stopped with 4% PFA.

### Quantitative RT-PCR

Total RNA was isolated from whole larvae with TRIzol reagent (Invitrogen, USA), and reverse transcribed to cDNA with HiScript III 1st Strand cDNA Synthesis Kit (+gDNA wiper) (Venzyme, Nanjing, China). Quantitative reverse-transcription PCR was performed using an Applied Biosystem 7500 Real-Time PCR system (Thermo Fisher, USA) with the ChamQ Universal SYBR qPCR Master Mix kit (Venzyme, Nanjing, China). The *eukaryotic translation elongation factor 1 alpha 1, like 1* (*eef1a1l1*) gene was used as the reference. Each of the biological replicates was run in 4 technical replicates. Relative gene expression was calculated by the following formula: Relative expression= 2^−(Ct[gene^ ^of^ ^interest]−Ct[housekeeping^ ^gene])^. Primer sequences are listed in supplementary Table 1.

### Heat-shock induction

Heat-shock induction was performed at various developmental stages as indicated in the text. For Heat-shock induction of *Tg(hsp70l:Tnfrsfa-mCherry)*, *Tg(hsp70l:DNtnfrsfa-mcherry)*, and *Tg(hsp70l:ikbSR-mcherry)* transgenes, larvae were placed in Holtfreter’s buffer and exposed to a 37°C water bath for 1 h, then reared at 28.5°C. The treatments were at 6 h intervals.

### Drug administration

For drug treatments, zebrafish larvae were placed in cell strainers in six well plates to facilitate transfer of larvae between solutions. All wells contained 10 mL of drug solution diluted with Holtfreter’s buffer. For hair cell ablation, 300 μM neomycin (Sangon, China) for 1 h; for Notch inhibition, 2 μM LY411575 (MCE, USA) from 1 dpf to 3 dpf. Holtfreter’s buffer with 1% DMSO was used for mock treatments. Following treatment, the fish were washed three times into fresh Holtfreter’s buffer by moving the cell strainers into adjacent wells.

### Statistical analysis

The posterior lateral line neuromasts (P1, P2, P3, and P4) from more than 7 zebrafish larvae were used for cell counting and measurement of the neuromast area. The area of neuromast was measured using Image J. The data were shown as Mean ± SEM. All statistical analyses were done with GraphPad Prism 6.0. The t-test was used for comparisons between two groups, whereas the Two-way ANOVA was used for determining whether there was an interaction effect between Notch signaling inhibition and *tnfrsfa* mutation on hair cell or support cell development. Statistical significance was set at p=0.05.

## Acknowledgements

This research was supported by the Intramural Research Program of the National Human Genome Research Institute (1ZIAHG200386-13).

## Supplementary Figures

**Figure S1.**
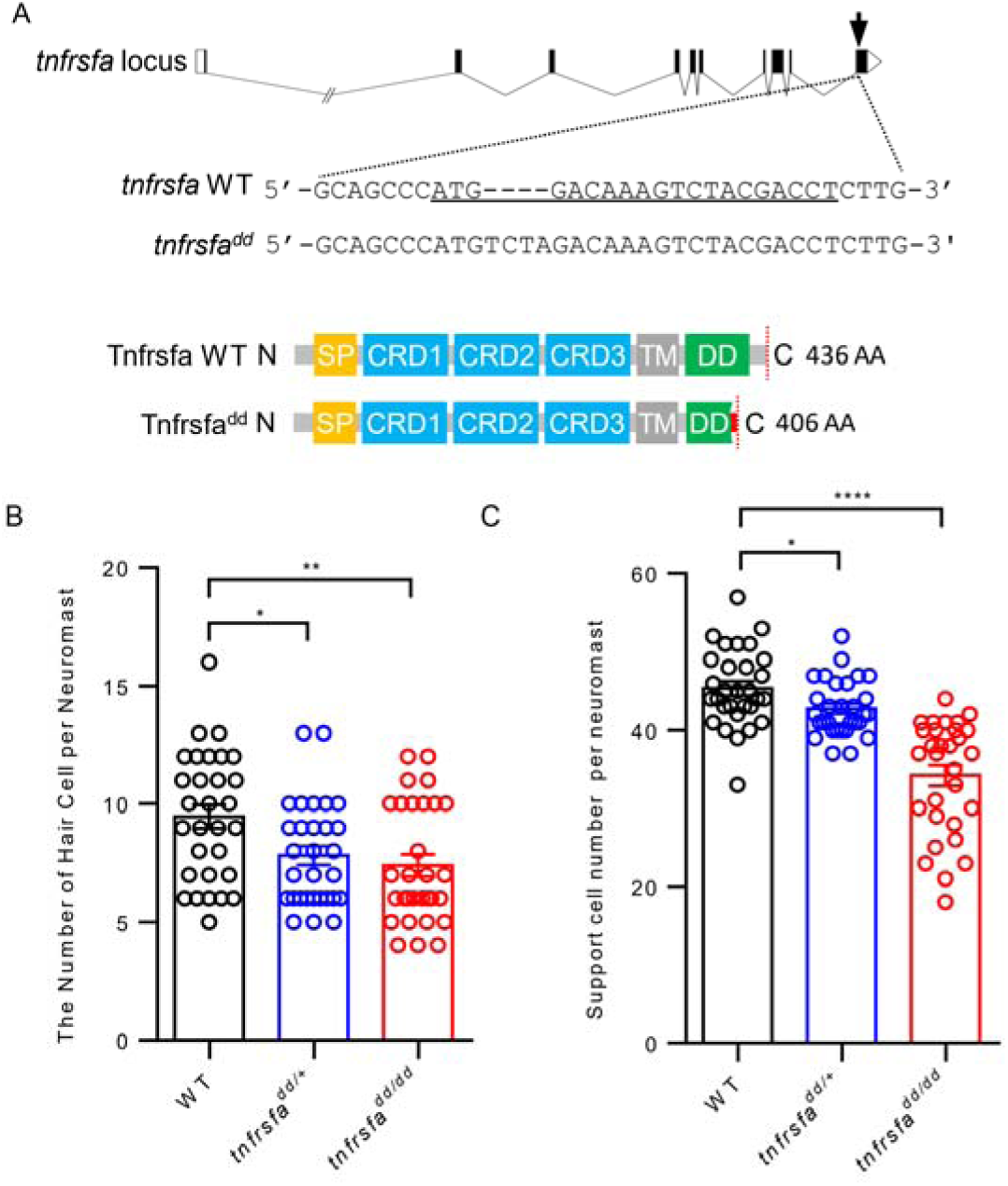
Disrupting the death domain in *tnfrsfa* reduces hair cells and accessory cells in neuromasts is sufficient to inactivate Tnfrsfa. Schematic diagrams showing the generation of *tnfrsfa* mutation by CRISPR/Cas9. Upper panel, the location of gRNA (arrow) in the *tnfrsfa* locus. Middle panel, the sequence surrounding the gRNA of wild-type and mutant *tnfrsfa*. The gRNA target is underlined. Bottom panel, the putative protein structure of wild-type and mutant Tnfrsfa. SP, signal peptide. CRD, cysteine-rich domain. TM, transmembrane domain. DD, death domain. Red box before the stop codon represents missense sequence. **B.** Quantification of hair cells in posterior lateral line neuromasts in wild-type, *tnfrsfa^dd/+^*, and *tnfrsfa^dd/dd^* larvae at 7 dpf using Yo-Pro-1 staining. Heterozygous mutants appear to have a dominant negative effect. C. Quantification of accessory cells in posterior lateral line neuromasts in wild-type, *tnfrsfa^dd/+^*, and *tnfrsfa^dd/dd^* larvae at 7 dpf by immunostaining with Sox2 antibody.

**Figure S2.**
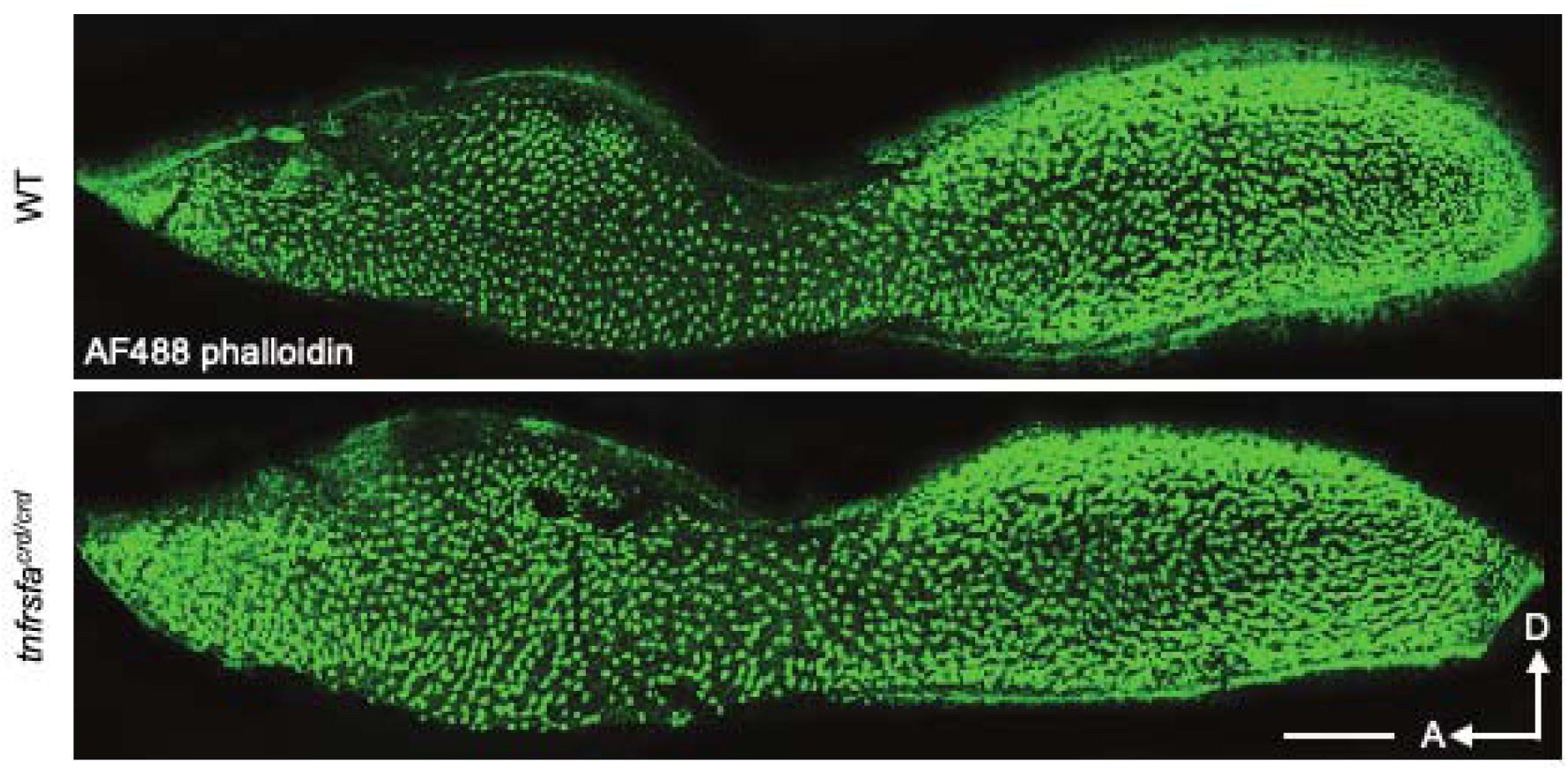
Inner ear development is normal in *tnfrsfa* mutants. Hair cells in the zebrafish inner ear saccule visualized using AF488 phalloidin in *tnfrsfa* mutant and wild-type sibling adults. No significant differences were detected. Scale bar, 100 μm. D, dorsal; A, anterior.

**Figure S3.**
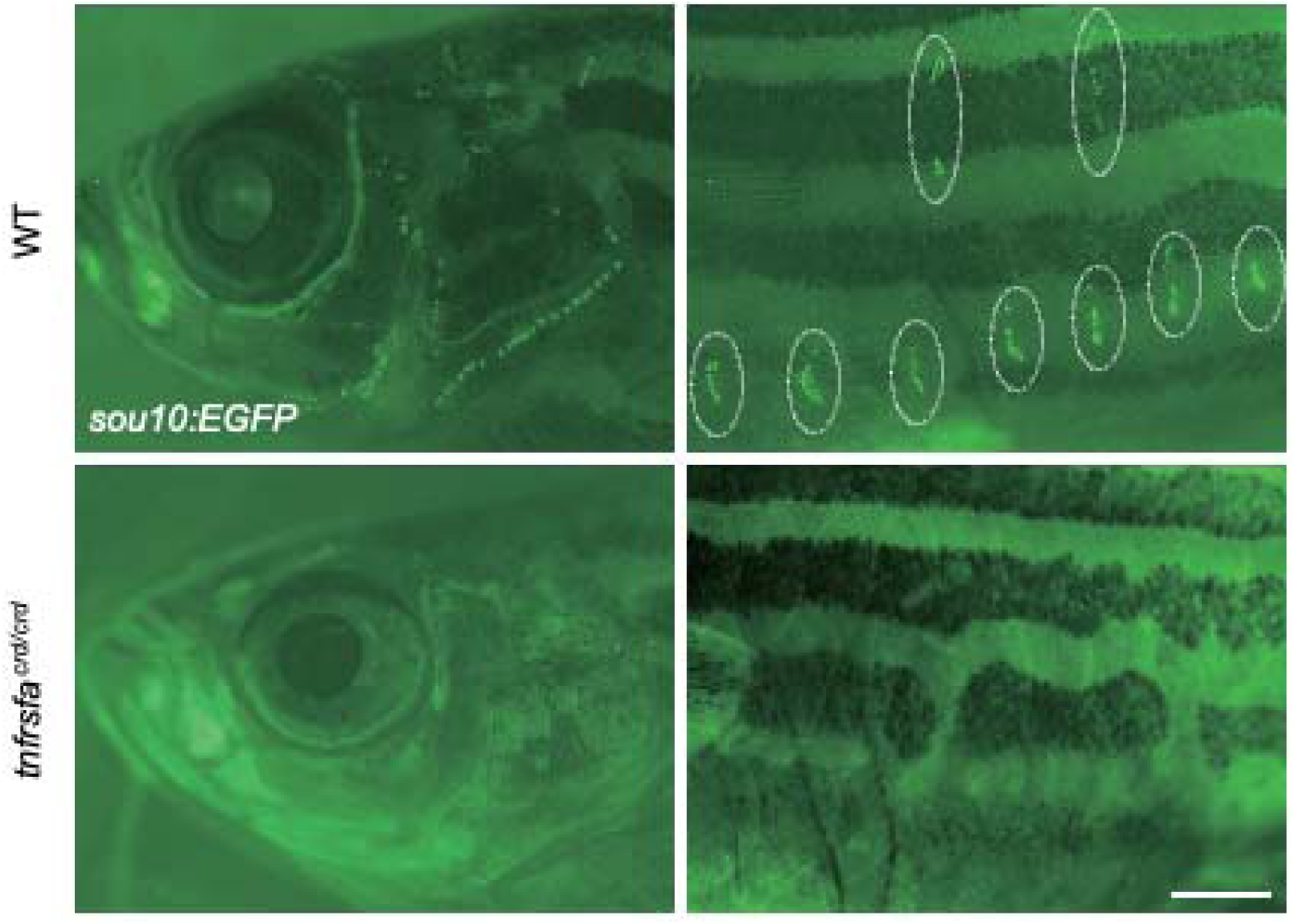
Adult lateral line neuromasts are missing in *tnfrsfa* mutants. Live imaging of support cells in *tnfrsfa^crd^* mutant and wild-type sibling adults at 3 mpf visualized by the *Tg(sou10:EGFP)* transgenic reporter line. White dashed circles outline superficial neuromast stitches. Scale bar, 100 μm.

**Figure S4.**
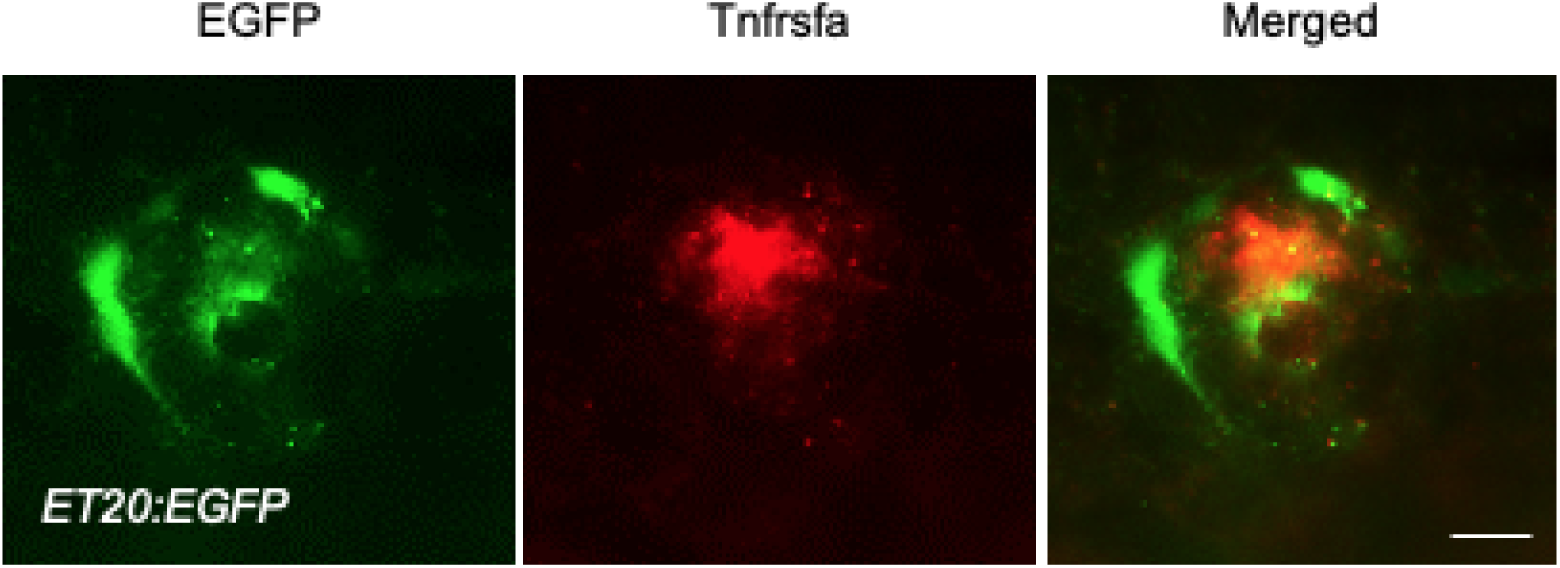
Tnfrsfa protein is strongly expressed in the dorsal support cells. Immunostaining of Tnfrsfa (red) and EGFP (green) in a *Tg(ET20:EGFP)* neuromast. No significant expression was detected in the GFP-positive cells. Scale bar, 10 μm.

**Figure S5.**
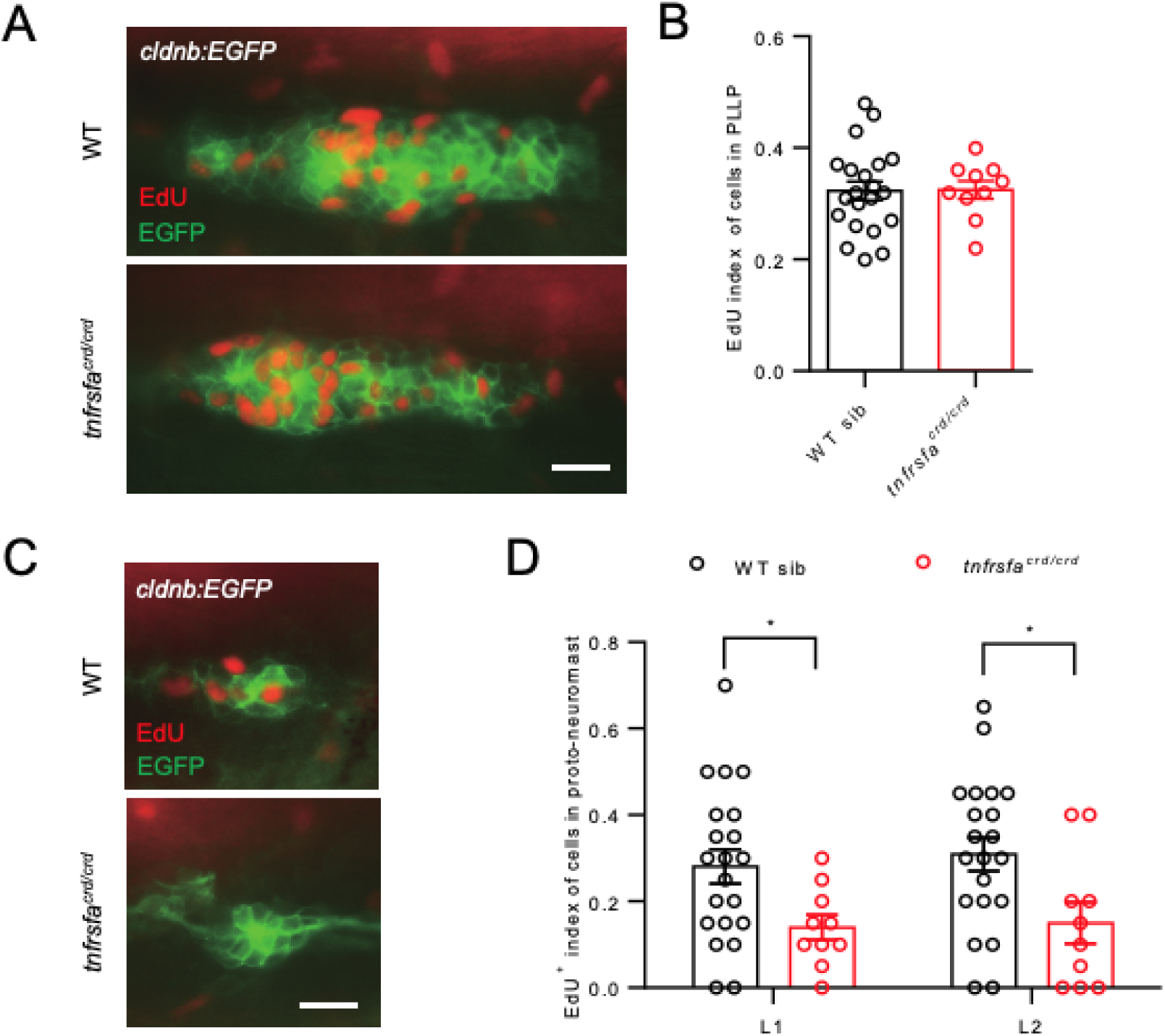
*tnfrsfa* plays a crucial role in cell proliferation in posterior lateral line proto-neuromasts. **A**, **C.** 2 h EdU incorporation in posterior lateral line primordium (**A**) and proto-neuromasts (**C**) visualized by *Tg(cldnb:EGFP)* at 30 hpf in wild-type and *tnfrsfa* mutant larvae. EGFP was detected by immunostaining with EGFP antibodies (green). Scale bar, 20 μm. **B, D**. Quantification of EdU indexes of posterior lateral line primordium (**B**) and proto-neuromasts (**D**) in wild-type and *tnfrsfa* mutant larvae in the L1 and L2 protoneuromasts. *, p < 0.05.

**Figure S6.**
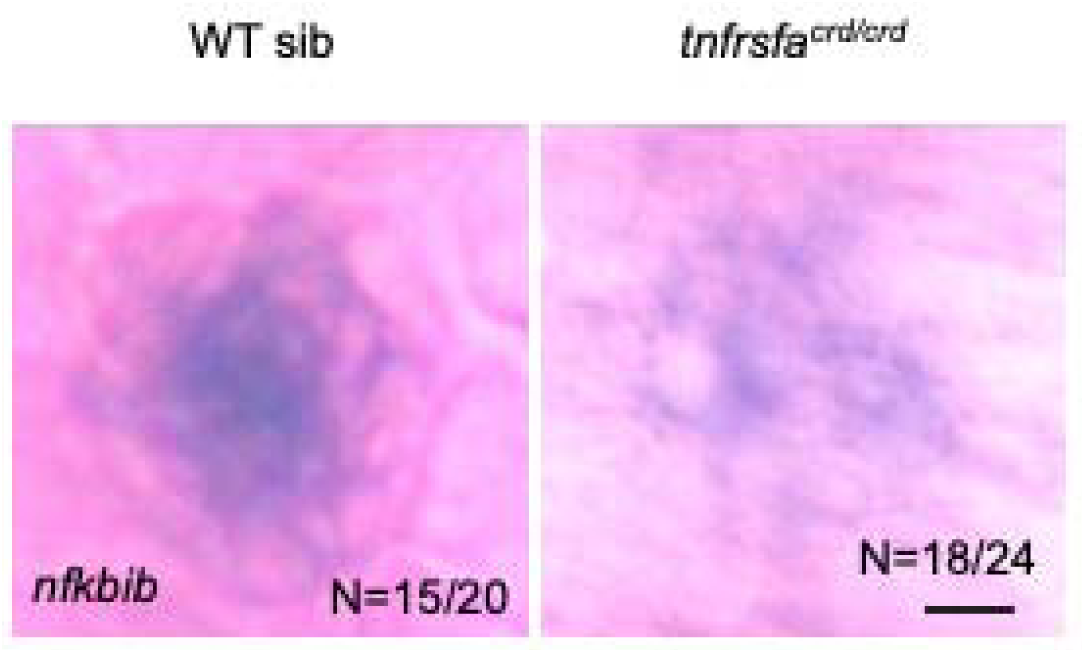
Tnfrsfa activates targets downstream of NF-κB signaling in the support cells. Whole-mount *in situ* hybridization of *nfkbib* in neuromasts of wild-type and *tnfrsfa* mutant larvae. Scale bar, 10 μm.

**Figure S7.**
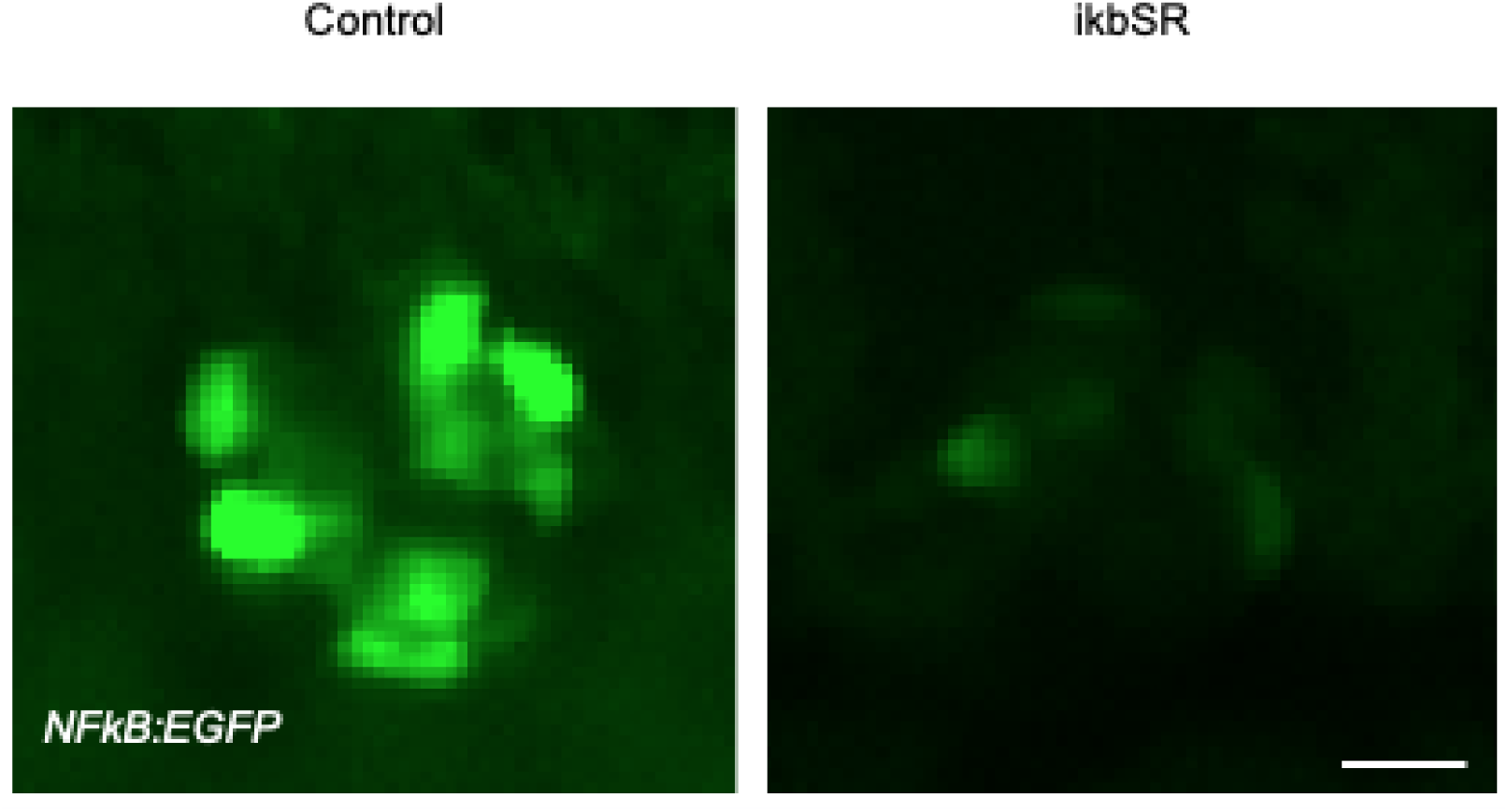
Overexpression of ikbSR inhibits NF-κB signaling in neuromast. Live imaging of neuromast from *Tg(NFkB:EGFP)* (Control) and *Tg(NFkB:EGFP)*; *Tg(hsp70l:ikbSR-mcherry)* (ikBSR) larvae at 5 dpf following heat-shock treatment for 2 days. Scale bar, 10 μm.

**Figure S8.**
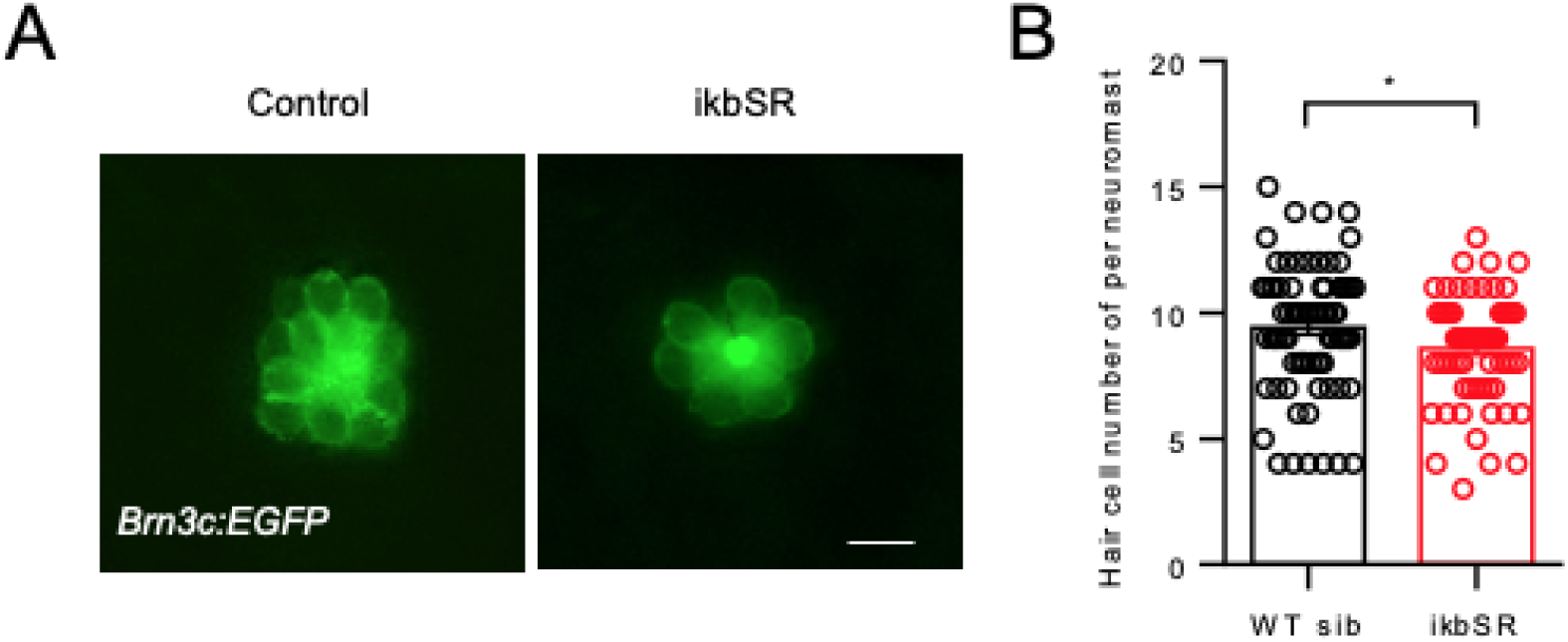
Overexpression of ikBSR reduces hair cells in posterior lateral line neuromasts. **A.** The live imaging of hair cells in neuromast from *Tg(Brn3:EGFP)* (Control) and *Tg(Brn3:EGFP)*; *Tg(hsp70l:ikbSR-mcherry)* (ikBSR). **B.** Quantification of hair cells in posterior lateral line neuromasts from *Tg(Brn3:EGFP)* (Control) and *Tg(Brn3:EGFP)*; *Tg(hsp70l:ikbSR-mcherry)* (ikBSR). Scale bar, 10 μm.

**Figure S9.**
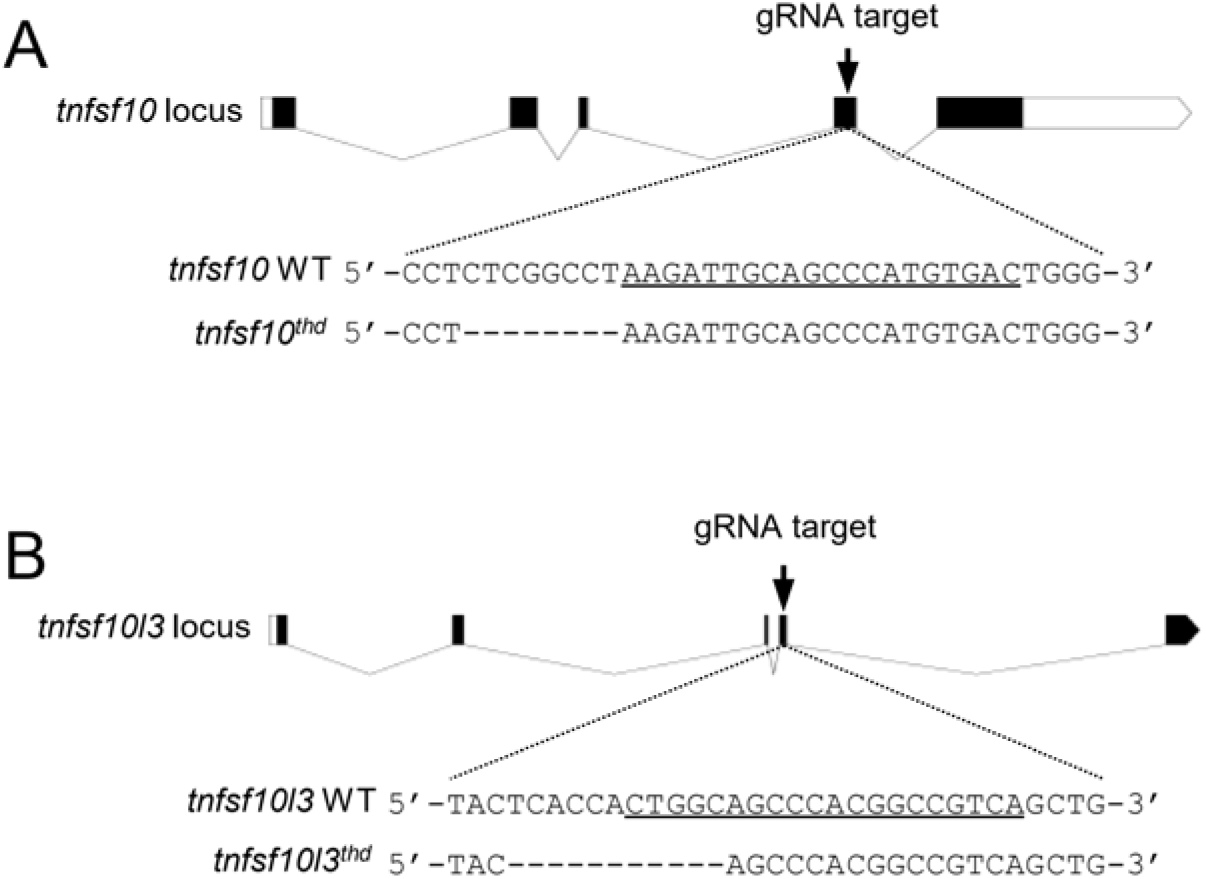
Targeted knockdown of the Tnfrsfa ligands. Schematic diagrams of generating *tnfsf10* (**A**) and *tnfsf10l3* (**B**) mutations generated by CRISPR/Cas9. Upper panel, the location of gRNA (arrow) in gene locus. Middle panel, the sequence around gRNA of wild-type and mutant. The gRNA target is underlined.

**Figure S10.**
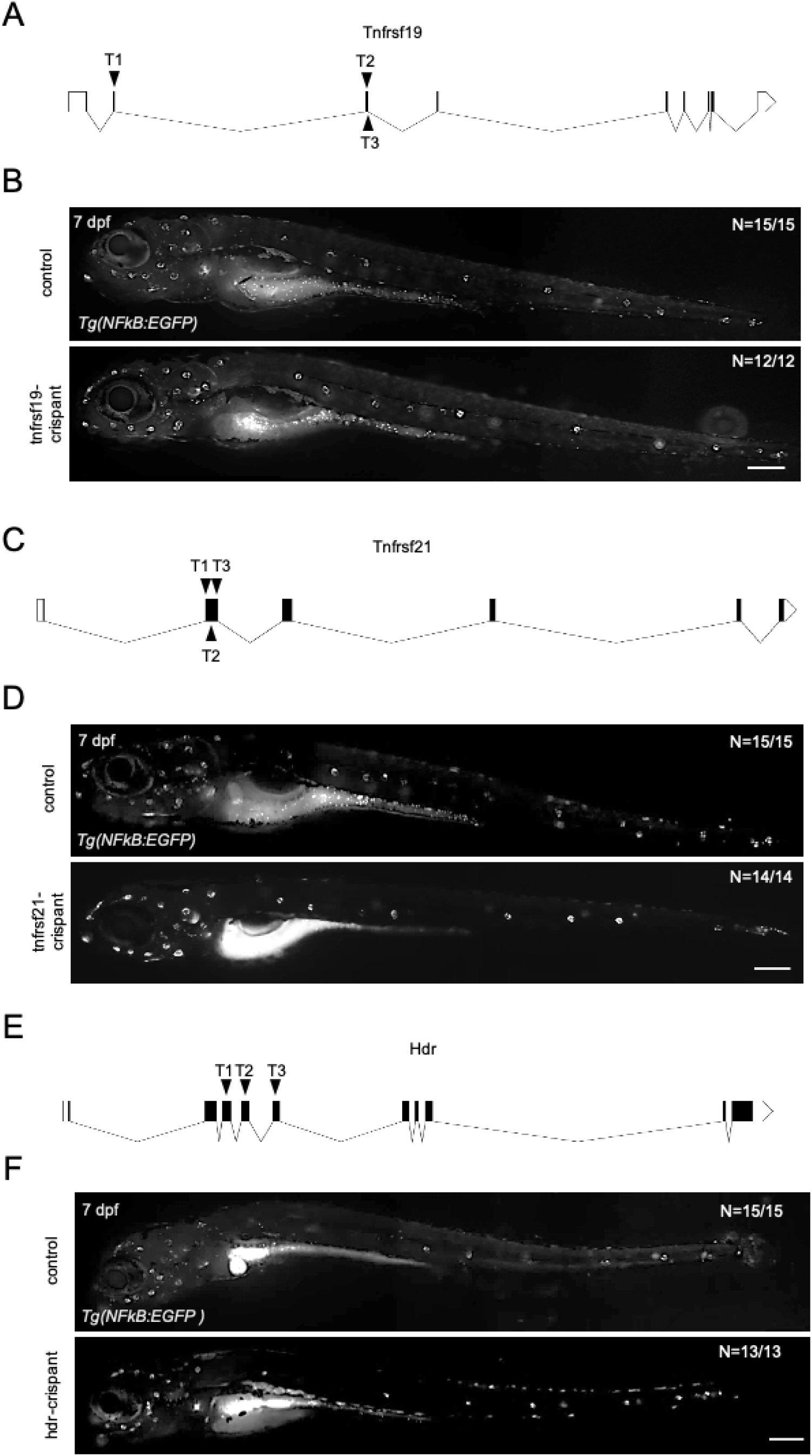
The impact of *tnfrsf19*, *tnfrsf21* or *hdr* mutations on NF-κB signaling in zebrafish lateral line neuromasts. A, **C**, and **E**: Schematic diagram of generating *tnfrsf19-, tnfrsf21-* and *hdr-crispants* using three gRNAs. **B**, **D**, and **F**: The fluorescence of *Tg(NFkB:EGFP)* in control and *tnfrsf19-, tnfrsf21-* and *hdr-crispants*. No major effects were seen. Scale bar: 200 μm.

**Figure S11.**
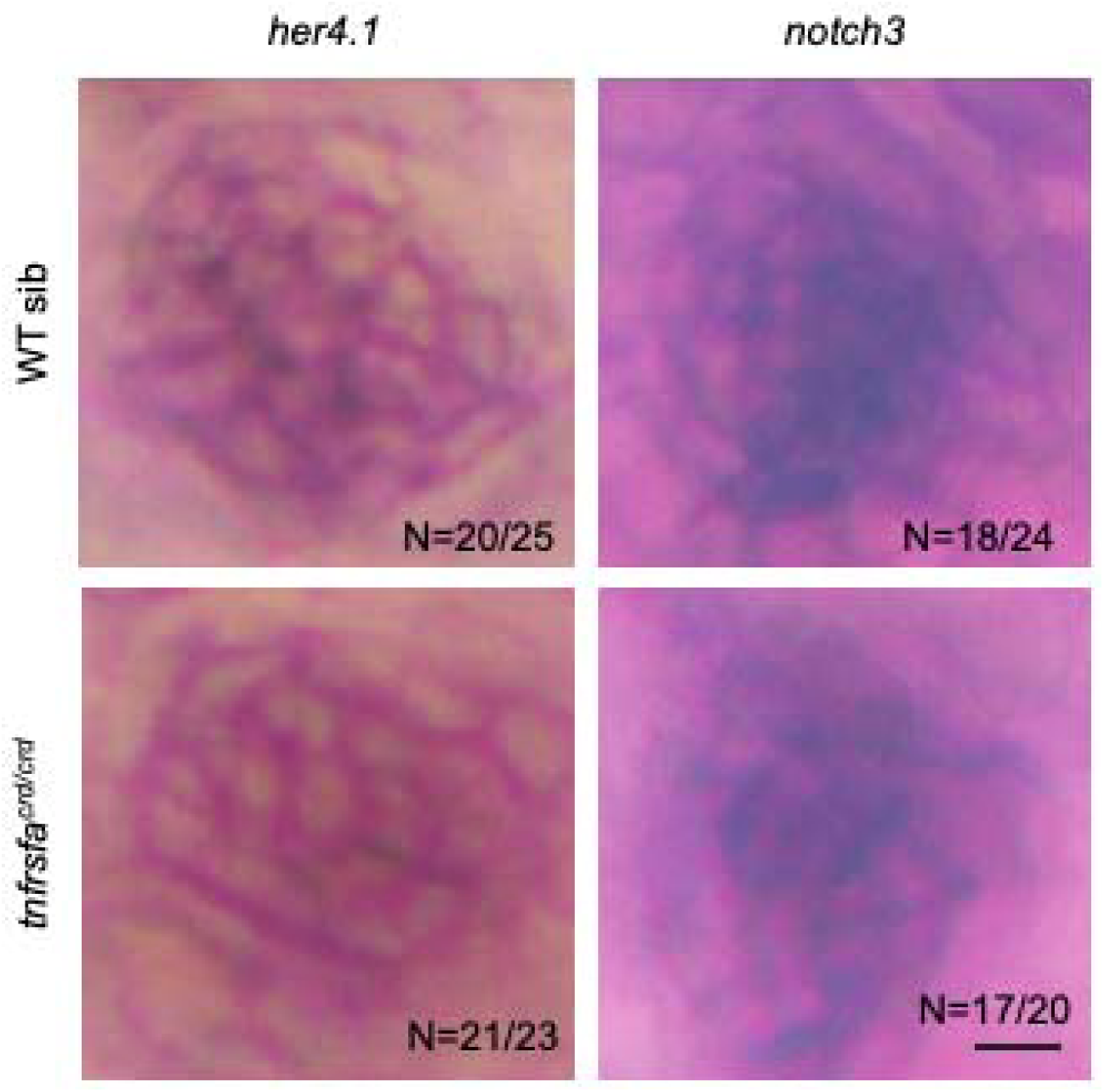
Mutations in tnfrsfa do not impact Notch activity. Whole-mount *in situ* hybridization showing the expression of *her4.1* and *notch3* in wild-type and *tnfrsfa* mutant zebrafish neuromasts. Scale bar, 10 μm.

